# Host-microbe interactions mediate doramectin-promoted metabolic reprogramming of CD8^+^ T-cells and amplify antitumor immunity

**DOI:** 10.1101/2023.01.29.525543

**Authors:** Sedigheh Taghinezhad-S, Amir Hossein Mohseni, Wen Jiang, Vincenzo Casolaro, Luis G. Bermúdez-Humarán, Florencia McAllister, Zhongwei Lv, Dan Li

## Abstract

The intestinal microbiota and its metabolites influence the host metabolic environment and CD8^**+**^ T-cell function. Metabolic changes in T-cells are thought to enhance the antitumor immune response. Here, we show that doramectin (DOR), a macrocyclic lactone (ML) of the avermectin (AVM) family, can modify CD8^+^ T-cell metabolism to increase and accelerate effector function. However, the functional capability of DOR depends mainly on the accessibility of gut microbiota. Using metagenomic and metabolomic techniques, we describe for the first time the interplay between gut microbiota and host metabolism involved in metabolic reprogramming of CD8^+^-T cells following DOR administration. Interestingly, we found that, after DOR administration, Firmicutes phylum not only impact DOR transport and absorption, but also boost amino acid levels in CD8^+^ T-cells, consistent with increased production of tumor necrosis factor alpha (TNF-α) and, in particular, interferon gamma (IFN-γ), which together play an important role in antitumor immunity. In contrast, the dysbiotic microbial community may abrogate the anticancer efficacy of DOR and lead to enhanced tumor growth and decreased survival. This finding likely supports the view that the presence of certain bacteria in the gut governs extra-intestinal immune responses and may be associated with metabolic adaptations necessary for efficient function of CD8^+^ T-cells upon DOR administration.

## Introduction

Several macrocyclic lactones (MLs) are members of the avermectin (AVM) family. The first development of AVM took place in the laboratories of the Kitasato Institute, following its isolation from a newly discovered soil-dwelling Actinomycetes, *Streptomyces avermitilis* (1). The AVM family includes a series of natural and semi-synthetic molecules, including ivermectin (IVM), abamectin (ABM), doramectin (DOR), eprinomectin, and selamectin, which are commonly used for the treatment of parasitic infections in different animal species (2). Members of the AVM family are extremely effective natural products that can be used as drugs in animals, humans, and crop protection (3). DOR [25-cyclohexyl-5-O-demethyl-25-de(1-methylpropyl)-avermectin A1a], a potent and novel IVM derivative macrocyclic lactone, has a closer structural similarity to ABM (4). In 1978, the U.S. Food and Drug Administration (FDA) approved the use of IVM, the most effective drug in the AVM family, in humans (5–7). Numerous studies have reported that IVM can be used to inhibit the proliferation of various types of cancer (8), such as ovarian cancer (9), renal cell carcinoma (10), hepatocellular carcinoma (11), melanoma (12), breast cancer (13, 14), lung cancer (11, 15), cervical cancer (16), gastric cancer (17), colorectal cancer (18), nasopharyngeal cancer (19), brain glioma (20), leukemia (21, 22), and human glioblastoma (23). In fact, IVM is recognized as an effective treatment for cancers resistant to chemotherapy drugs (24, 25).

Despite these exciting findings, the impact of DOR in cancer treatment remains largely unknown. We hypothesized that DOR may serve as a novel therapeutic candidate to fight cancer. The main objectives of this study were to explore DOR impact on CD8^+^ T-cell stimulation and its ability to extend survival in animal models. We found that DOR promoted CD8^+^ T-cell-mediated antitumor immunity *in vitro* and *in vivo*. In particular, we demonstrate that metabolic and functional alterations in CD8^+^ T-cells subsequent to DOR administration, partly mediated by the gut microbiota and its metabolites, could potentially enhance the response to chemotherapy, offering novel evidence of crosstalk between commensal microbiota and the host immune system. Thus, the metabolic host-microbe-DOR interactions discovered in the present study describe a new mechanism of bacterial-mediated drug metabolism, paving the way for future studies exploring the impact of microbiota on the DOR-mediated antitumor effect in different mammalian systems and their possible translation into clinical trials.

## Results

### 1. *In vitro* effect of DOR on CD8^+^ T-cell activation

We first determined whether DOR impacts antitumor immune functions by assessing its effects on CD8^+^ T-cell responses. Strikingly, the addition of DOR significantly potentiated mouse and human CD8^+^ T-cell activation triggered by anti-CD3ε and -CD28 antibodies, with 2.5 μM being its maximally effective concentration (Figure 1A-C). Following inflammatory and antigenspecific signals, naïve T-cells soon undergo massive expansion including rapid division and differentiation. Notably, DOR treatment resulted in a significant increase in the percentages of naïve CD8^+^ T-cells as well as CD62L^high^CD44^high^CD8^+^, central memory T-cells, but not CD62L^low^CD44^high^CD8^+^, effector memory T-cells, in spleen and mesenteric lymph nodes (MLNs) (Figure 1D). Similarly, treatment of mouse and human CD8^+^ T-cells with DOR, compared with DMSO (vehicle control), increased activation, and expansion of TNF-α^+^CD8^+^ and IFN-γ^+^CD8^+^ T-cells in a dose-dependent manner (Figure 2A-C). Moreover, DOR, but not DMSO alone, induced a substantial increase in activation of OT-I mouse CD8^+^ T-cells stimulated by ovalbumin (OVA)_257-26_4(SIINFEKL) peptide. Consistent with this, DOR supplementation led to an increased percentage of CD8^+^ T-cells producing tumor necrosis factor α (TNF-α) and interferon gamma (IFN-γ), but not granzyme B (Figure S1A). Co-culture with OT-I CD8^+^ T-cells activated in the presence of DOR triggered increased apoptosis of B16-OVA cells (Figure S1B, left) and human papillary thyroid carcinoma BCPAP cells (Figure S1B, right). Activation of the Src family tyrosine kinase LCK is required to initiate antigen-specific TCR signaling in T-cells. Notably, when CD8^+^ T-cells were supplied with DOR, relative to DMSO alone, the phosphorylation levels of LAT (Tyr255), LCK (Tyr394), PI3K (Tyr458), and Akt (Ser473) were significantly increased following activation (Figure 2D). In addition, we observed metabolic reprogramming of DOR-treated CD8^+^ T-cells after activation. In particular, the levels of 12-hydroxydodecanoic acid, methyl jasmonate, 3-hydroxy capricacid, N-methyl-L-glutamic acid, and fexofenadine were remarkably increased in these cells, whereas the levels of isokobusone, cholesterol sulfate, ketoleucine, and tridecanoic acid were markedly decreased (Figure 2E). This suggests that the enhancement by DOR of T-cell activation is paralleled by metabolic reprogramming.

**Figure 1:**
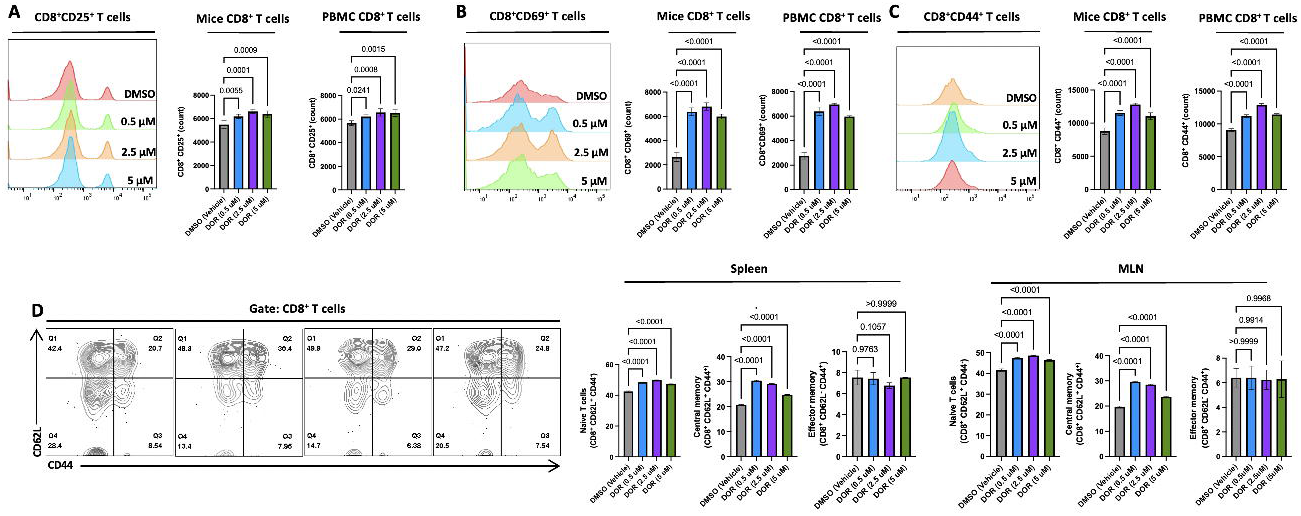
Identification of DOR as a promotor of CD8^+^ T-cell activation. **A-C:** naive mouse or human CD8^+^ T-cells were activated by anti-CD3ε and -CD28 antibodies in complete medium containing 0.5, 2.5, or 5 μM DOR or DMSO (vehicle control) for 24 h (to detect CD69 expression, B) or 40 h (to detect CD25 and CD44 expression, A and C). **D:** Coexpression of CD62L and CD44 on the surface of different subpopulations of CD8^+^ T-cells from mouse spleen (center) and MLN (right). Representative flow cytometry plots are shown with the percentages of cells indicated in each quadrant (left). All data are the mean ± SEM of n = 3 independent wells per experiment.

**Figure 2:**
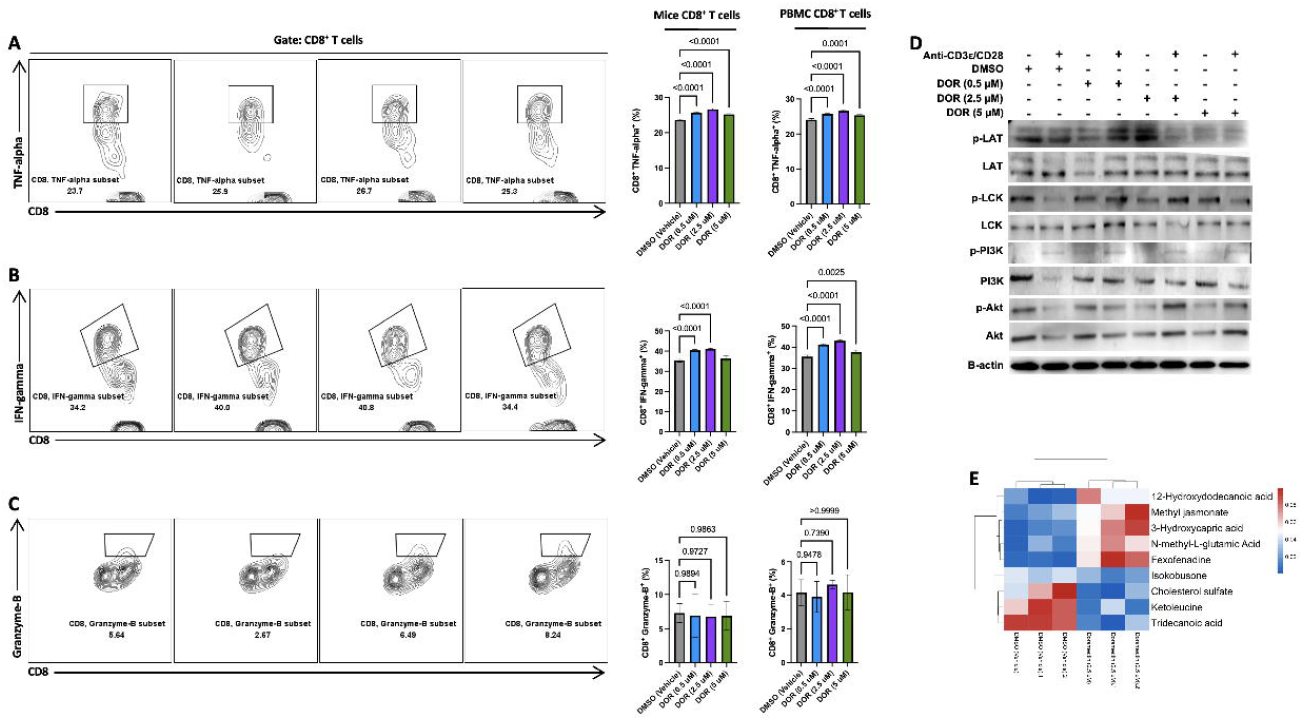
**A-C:** Naive mouse or human CD8^+^ T-cells stimulated by anti-CD3ε and -CD28 antibodies in complete medium containing 0.5, 2.5, or 5 μM DOR or DMSO (vehicle control) for 4 h were analyzed for expression of the TNF-α^+^ (A), IFN-γ^+^ (B), and Granzyme B^+^ (C) using FACS. **D:** Mouse CD8^+^ T-cells left unstimulated (naive) or stimulated (5 μg mL^−1^ anti-CD3ε and 1 μg mL^−1^ anti-CD28 antibodies) for 15 min in the presence of increasing quantities of DOR or DMSO (vehicle control) were analyzed for expression of the indicated proteins by Western blotting. **E:** Mouse naive CD8^+^ T-cells were activated in medium containing DOR (2.5 μM) or DMSO (vehicle control) prior to stimulation with 5 μg mL^−1^ plate-coated anti-CD3ε and 1 μg mL^−1^ soluble anti-CD28 (1.5 h at 37°C and 5% CO_2_). All media were added with 10% dialysed FBS. The abundance of cellular metabolites was analyzed by LC-MS/MS. The heat map shows the relative abundance of metabolites in CD8^+^ T-cells stimulated in DOR-supplemented relative to control medium. Data are representative of three independent experiments.

### 2. *In vivo* effect of DOR on CD8^+^ T-cell activation

To evaluate the impact of DOR on the endogenous antitumor immune response, we intraperitoneally (i.p.) injected 200 μg/kg (200 N group) and/or 350 μg/kg (350 N group) of DOR, or PBS vehicle control (V N group) into wild-type (WT) C57BL/6 mice, starting 7 days after subcutaneous (s.c.) inoculation of melanoma tumor cells (B16-F10, 10^6^ cells/mouse), then every second day through the remainder of the experiment (Figure 3A). Mice injected with either 200 μg/kg or 350 μg/kg DOR developed smaller tumors (Figure 3B, up) and exhibited reduced spleen size (Figure 3B, bottom). However, 200 N mice not only exhibited reduced tumor size and retarded growth of melanoma cells (Figure 3C), but also diminished weight loss compared with V N or 350 N mice (Figure 3D). In addition, the group of mice that was treated with 200 μg/kg DOR (group 200 N) displayed the most increased survival (Figure 3E). Flow cytometry data showed no significant changes were seen in the frequencies of CD8^+^CD25^+^ T-cells (Figure 3F) and memory CD8^+^ T-cells (Figure 3G) in the Peyer’s patches (PPs) from either DOR-injected group compared to the V N group. In contrast, a significant increase in the frequencies of CD8^+^CD25^+^ T-cells (Figure 3H), naïve T-cells (Figure 3I, left), and central memory CD8^+^ T-cells (Figure 3I, center), but not effector memory T-cells (Figure 3I, right) in MLN from either DOR-injected group. Antibody-mediated depletion of CD4^+^ and CD8^+^ T-cells in DOR-treated mice reduced tumor growth suppression (Figure S1C), pointing to a pivotal role for both CD4^+^ and CD8^+^ T-cells in DOR-promoted antitumor immunity.

**Figure 3:**
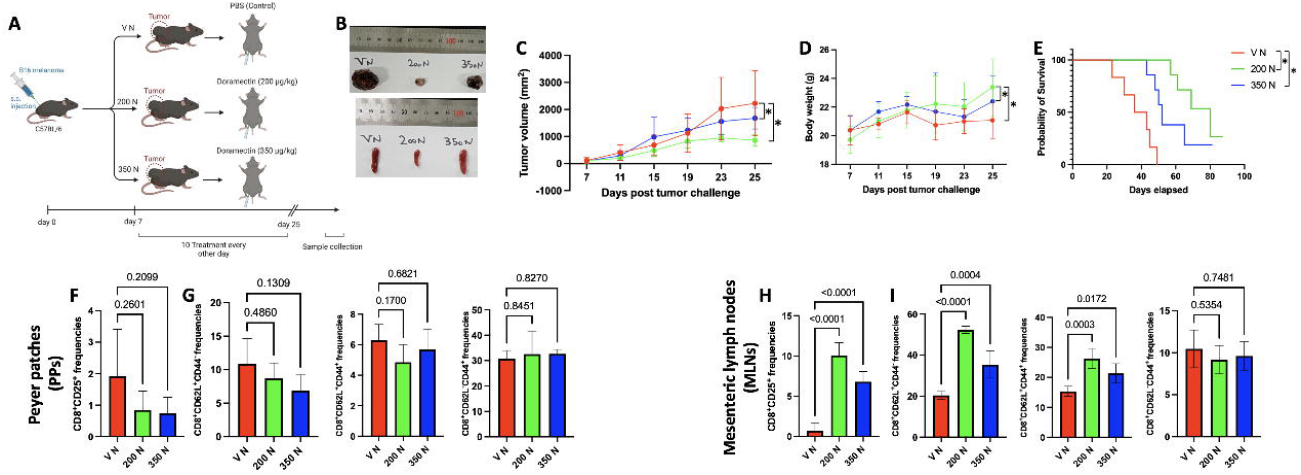
**A-E:** C57BL/6 mice were s.c. transferred with 10^6^ B16-OVA cells and, after 7 days, i.p. injected with the indicated concentrations of DOR (200 or 350 μg/kg), or PBS containing corresponding amounts of DMSO (vehicle control) (A). After 25 days the tumors and spleens were explanted, and their representative images are shown (B). The tumor burden (C), body weight (D), and survival (E) over time were also assessed as indicated. Asterisks are representative of p-values < 0.05. **F-I:** DOR enhances CD8^+^ T-cell activity and responses *in vivo*. Percentages of CD25^+^CD8^+^ T-cells were measured in the Peyer patches (PPs; F) and mesenteric lymph nodes (MLNs; H). Also measured were the percentages of CD62L^+^CD44^-^ CD8^+^ T-cells (G and I, left), central (CD62L^+^CD44^+^CD8^+^ T cells; G and I, center) and effector (CD62L^-^CD44^+^CD8^+^; G and I, right) memory-like CD8^+^ T-cells in the PPs and MLN. All data are the mean ± SEM of n = 3 independent determinations.

Independent evidence in support for T-cell activation in DOR-treated mice was obtained in experiments using a transgenic mouse model (Figure S1D). After s.c. injection of murine B16/OVA melanoma cells (H2b) for 7 days into the dorsal region of C57BL/6 mice expressing the CD45.2 allele, mice were immunodepleted by irradiation (2 Gy, twice), then adoptively transferred with 5 × 10^5^ OT-I TCR transgenic T-cells expressing the CD45.1 allele and primed by OVA_257–264_ peptide (SIINFEKL). Mice were i.p. injected with PBS containing 200 μg/kg DOR (200 N) or a corresponding amount of DMSO (V N), starting one day before injection of OVA_257–264_ peptide, then every second day through the remainder of the experiment (Figure S1D). The data are in line with the idea that OT-I CD8^+^ T-cells were able to mount an effective antitumor response after DOR administration (Figures S1E-F). Higher frequencies of OT-I CD8^+^CD45.1^+^ central memory T-cells, not effector memory OT-I CD8^+^CD45.1^+^ cells, in the spleen (Figure S1G) suggest either enhanced recruitment or higher survival of OT-1 T-cells in DOR-treated mice.

In subsequent experiments, we studied the mechanism(s) underlying T-cell activation by DOR in this model. Our data showed that activation of OT-I CD8^+^ T-cells increased phosphorylation of mTORC1 (Ser151) and S6 (Ser235/236) (Figure S1H) in the spleens of 200 N mice when supplied with DOR, and this effect was potently intensified by rapamycin supplementation (Figure S1I). Taken together, these results indicate a previously unrecognized role of DOR in the regulation of TCR signaling.

### 3. Gut microbiota shapes antitumor immunity by modulating DOR efficacy

Increasing evidence supports the critical role of the gut microbiota in the control of immune surveillance and its ability to modulate host immune responses to therapy (26, 27). Therefore, we wondered whether differences in gut microbiota composition might modify the immune phenotypes associated with tumor growth inhibition in DOR-treated mice. To test this hypothesis, mice received an antibiotic cocktail (ABX) in the drinking water 5 days prior to tumor inoculation (Figure 4A). Strikingly, ABX treatment largely impaired the ability of DOR to interfere with melanoma growth in mice treated with either 200 (200 A) or 350 μg/kg DOR (350 A) (Figure 4B), suggesting that the gut microbiota plays a key role in the ability of DOR to control tumor growth in this model. In particular, the altered gut microbiota profiles in these mice were associated with impaired tumor growth inhibition (Figure 4C), increased weight loss (Figure 4D), and decreased survival (Figure 4E). Thus, we concluded that administration of DOR to ABX-treated mice elicited immune perturbations in PPs and MLN in response to gut dysbiosis (Figure S2A-H). Moreover, germ-free (GF) animals, compared to specific pathogen free (SPF) mice, similarly failed to respond to DOR treatment (Figure 4F). These observations point to the importance of the gut microbiota in the control of antitumor immunity following DOR administration.

**Figure 4:**
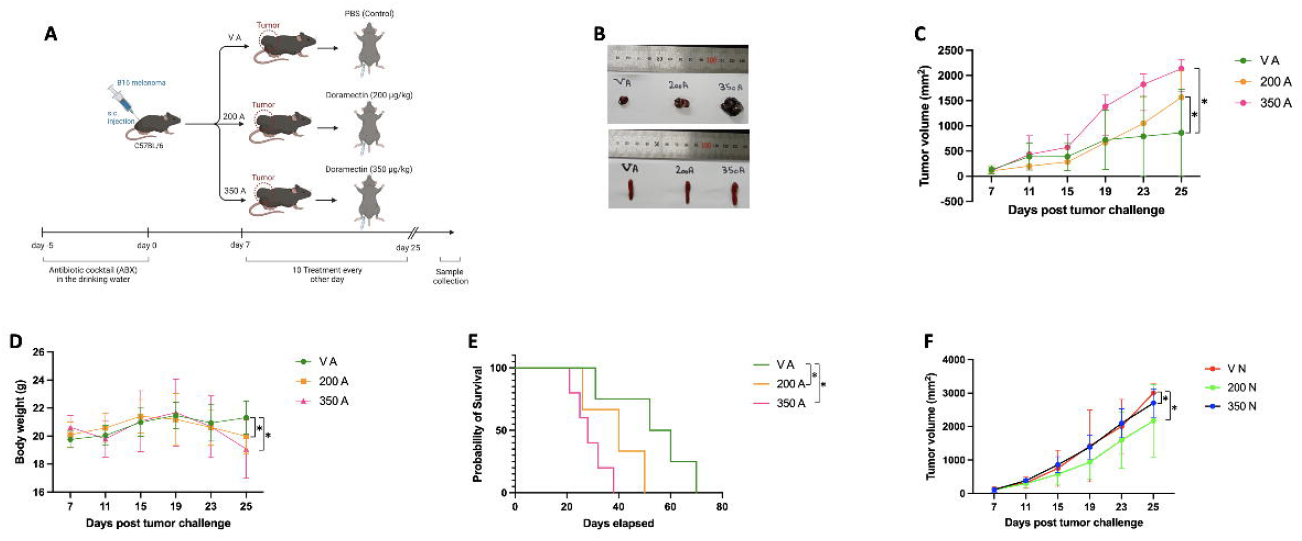
**A:** C57BL/6 mice were treated with ABX for 5 days prior to tumor inoculation then injected with 200 or 350 μg/kg of DOR, or PBS containing equal amounts of DMSO (vehicle control). **B-E:** Tumor and spleen sizes (B), tumor volume (C), body weight (D), and mice survival (E) were assessed. **F:** Germ-free C57BL/6 mice were s.c. injected with 10^6^ B16-OVA cells and after 7 days were i.p. injected with 200 or 350 μg/kg of DOR, or PBS containing equal amounts of DMSO (vehicle control). Shown is the tumor burden over time. Data are the means ± SEM of n = 3 independent determinations. Asterisks are representative of p-values < 0.05.

### 4. The phylum Firmicutes mediated the DOR effect

We then explored whether the immune phenotypes observed in the normal mice groups (200 N and 350 N, Figure 3A) *versus* the antibiotic-treated mice groups (200 A and 350 A, Figure 4A) were related to, or affected by, specific alterations in the composition of the gut microbiota. To map gut microbiota composition, we used 16S rRNA amplicon sequencing of V3-V4 regions followed by computational analysis to profile the fecal microbiota from all groups 18 days after DOR injection (corresponding to 25 days after tumor cell inoculation). According to principal component analysis (PCA), the microbial communities of the 200 N group formed a separate cluster from other groups (Figure 5A). These data confirmed that DOR significantly reshaped the composition of the microbiota of the small/large intestine in 200 N mice bearing s.c. tumors, resulting in a significant decrease of bacterial communities of the phylum Firmicutes, distributed across 12 species, when compared with the V N group (Figure 5B, Figure S3A, and Figure S3B). Among them, eight phylotypes (Figure 5C), such as *Lachnoclostridium*_sp._An169, *Dorea_sp._5-2, Eubacterium plexicaudatum, Lachnospiraceae* bacterium 3-1, *Eubacterium* sp. 14-2, uncultured *Clostridium* sp., *Anaerotruncus* sp. G3(2012), and *Lachnospiraceae* bacterium A2 were abundant in V N group, while ABX administration led to their elimination (Figure 5C). We inferred then that these eight phylotypes positively correlated with decreased tumor size (Figure 3C) and increased survival (Figure 3E) in 200 N mice. In contrast, no significant alterations were observed in the composition of gut microbiota in the 200 A and 350 A groups compared to V A (Figure 5B and Figure S3A). Notably, colon PPs CD8^+^IFN-γ^+^ T-cells (Figure 5E) and Granzyme B^+^ T-cells (Figure 5F), but not TNF-α^+^ (Figure 5D), were reduced in 200 N mice compared to the V N group, whereas a concomitant, significant increase of CD8^+^TNF-α^+^ (Figure 5G) and CD8^+^IFN-γ^+^ T-cells (Figure 5H), but not Granzyme B^+^ cells (Figure 5I), was observed in the MLN, proposing that bacterial translocation upon administration of DOR contributed to the reduced frequencies of activated T-cells (Figure 3F) and CD8^+^IFN-γ^+^ T-cells (Figure 5E) in the gut. Vice versa, our results showed no significant increase in the levels of activated T-cells (Figure S2E), CD8^+^TNF-α^+^ (Figure S2F), and CD8^+^IFN-γ^+^ T-cells (Figure S2G) in the MLN from groups 200 A and 350 A relative to levels found in their PPs (Figures S2A-C), highlighting the failure of DOR in promoting bacterial translocation in the presence of dysbiosis in the gut microbial communities. To confirm the contribution of bacterial translocation, we sterile collected and dissociated MLN and spleen in an aerobic or anaerobic chamber. After 48 h of incubation, we observed significant translocation of commensal bacteria into the MLN and spleen after DOR treatment in the normal mice groups (200 N and 350 N) (Figure S2I, K, M, and O), but not in the corresponding ABX groups (200 A and 350 A) (Figure S2J, L, N, and P).

**Figure 5:**
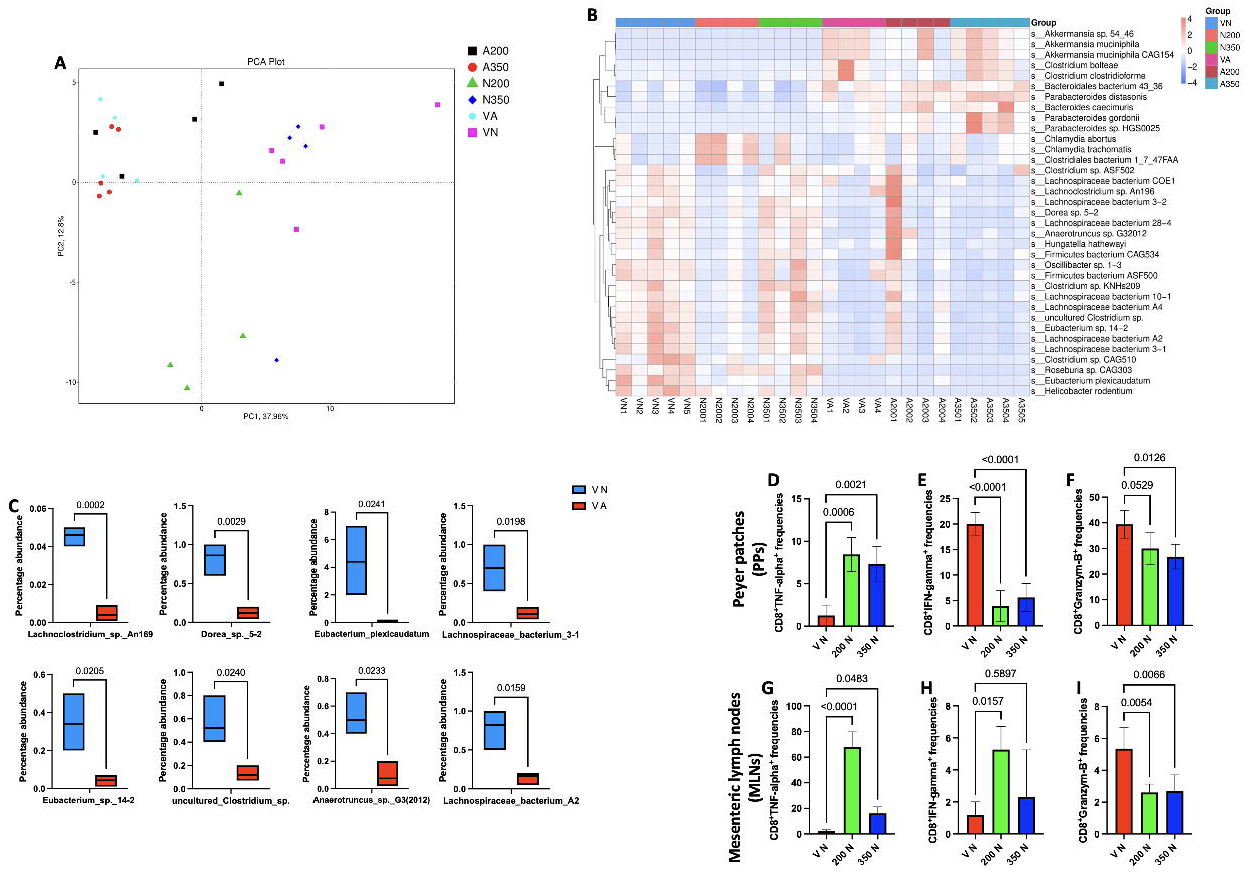
**A:** Scatter plot for PCA model for normal groups (V N, 200 N, and 350 N) as well as ABX-treated groups (V A, 200 A, and 350 A). The abscissa PC1 and the ordinate PC2 in the figure represent the scores of the first and second principal components, respectively, each scatter point represents a sample, and the color and shape of the scatter point represent different groups. The sample is basically within the 95% confidence interval (Hotelling’s T-squared ellipse). **B:** Abundance clustering heatmap based on significantly different species. Horizontal is sample information; vertical is species annotation information; on the left side of the figure is the species clustering tree; the value corresponding to the middle heatmap is the Z-value obtained after the relative abundance of the species row is normalized. **C:** Boxplot diagram of the relative abundance of the eight dominating phylotypes in the V N group compared to V A. **D-I:** Percentages of CD8^+^TNF-α^+^ (D and G), CD8^+^IFN-γ^+^ (E and H), and Granzyme B^+^CD8^+^ T-cells (F and I) were measured in the PPs and MLNs in groups V N, 200 N, and 350 N by FACS. Differences were analyzed with the Student’s *t*-test. All data are the mean ± SEM of n = 3 independent determinations.

Furthermore, in the 200 N group, compared to V N, the relative abundance of 3 species, which mapped most closely with *Clostridiales* bacterium 1-7-47FAA, *Chlamydia abortus*, and *Chlamydia trachomatis*, was predominantly increased (Figure S3C), confirming that a decreased level of activated T-cells (Figure 3F) and CD8^+^IFN-γ^+^ T-cells (Figure 5E) in the gut could be accounted for by bacterial translocation. However, none of these species were positively related to tumor size, weight loss, and survival. Altogether, these results highlight the ability of DOR to induce selective translocation of different Gram-positive bacterial species in the presence of significant alterations in the small/large intestinal microbiota (Figure 3C-E).

We identified 5 phylotypes that were robustly enriched after ABX administration in group V A relative to V N. Of these, *Akkermansia* sp.-54-46, *A. muciniphila*, and *A. muciniphila* CAG:154, were significantly increased in group V A (Figure S3D), suggesting that, in the absence of DOR, short-term ABX administration led to an increase of *A. muciniphila* species, possibly accounting for decreased tumor size (Figure 4C) and, in turn, increased survival in this group (Figure 4E) (28).

### 5. Metabolic changes in the gut after DOR administration

To explore the metabolic and functional effects of microbial community changes across the different treatment groups, we used evolutionary genealogy of genes: Non-supervised Orthologous Groups (eggNOG), Carbohydrate-Active enZymes (CAZy) and Kyoto Encyclopedia of Genes and Genomes (KEGG) annotation programs to get the full breadth of functions in each group. We identified several hundred differentially expressed genes between the 200 N and V N groups as well as between the 200 A and V A groups (Table S1-2). These data showed that DOR administration led to a substantial decrease in the representation of the KEGG categories of 6 main processes (Figure 6A), 24 eggNOG orthologues main processes (Figure S4A) and 6 CAZy main processes (Figure S5A) in the 200 N group. For example, the relative abundance of genes encoded in KEGG level 2 pathways shifted so that carbohydrate metabolism, amino acid metabolism, translation, and membrane transport among others were the most significantly diminished among the microbiota of the 200 N group (data not shown). In addition, PCA analysis revealed that the 200 N group tends to cluster separately from other groups, implying that this sample has high similarity in community structure (Figure 6B, Figure S4B, and Figure S5B). The abundance clustering heatmap based on significantly different functions indicates that the 200 N group is extremely different from the other groups (Figure 6C, Figures S4C, and Figures S5C). Furthermore, we speculate that the distinctive antitumor cellular immune response in this group is related to DOR and, particularly, with the metabolic processes involved in DOR uptake by the cells. To verify this hypothesis, we used the Metastats method, and a significant difference function box plot was drawn based on the q-value. We found that 9 KOs, including K03088, K01990, K02003, K07484, K02004, K06147, K03205, K03496, and K00558, were significantly decreased in the fecal microbiota of the 200 N compared with the V N groups (q-value < 0.05). Next, we aimed to study whether the decreased orthologs matched with certain physiological processes. For this, we examined the overrepresentation of these orthologs in KEGG pathways. Our analysis showed several intriguing associations. Specifically, among 9 KOs that were significantly decreased in the fecal microbiota of 200 N compared with V N mice, we found that the microbiota of the V A group *versus* that of V N mice was significantly decreased in orthologs correlated with ABC type transport system ATP-binding proteins, such as K01990 (ABC-2 type transport system ATP-binding protein) (Figure 6D), K02003 (putative ABC transport system ATP-binding protein) (Figure 6E), and K02004 (putative ABC transport system permease protein) (Figure 6F), emphasizing the significance of these pathways for metabolizing DOR. We therefore reasoned that dysbiotic events in gut microbiota and the subsequent loss of these ABC transporters upon eradication of specific gut microbiota after ABX administration result in increased susceptibility to low (200 μg/kg) and, more so, high DOR concentrations (350 μg/kg), thereby leading to DOR accumulation and generating toxic consequences, which account for increased tumorigenesis (Figure 4C) and low survival (Figure 4E). Overall, these results unravel important metabolic activities and potential functional Firmicutes-host interactions that mediate DOR metabolism and inherently CD8^+^ T-cell immunity.

**Figure 6:**
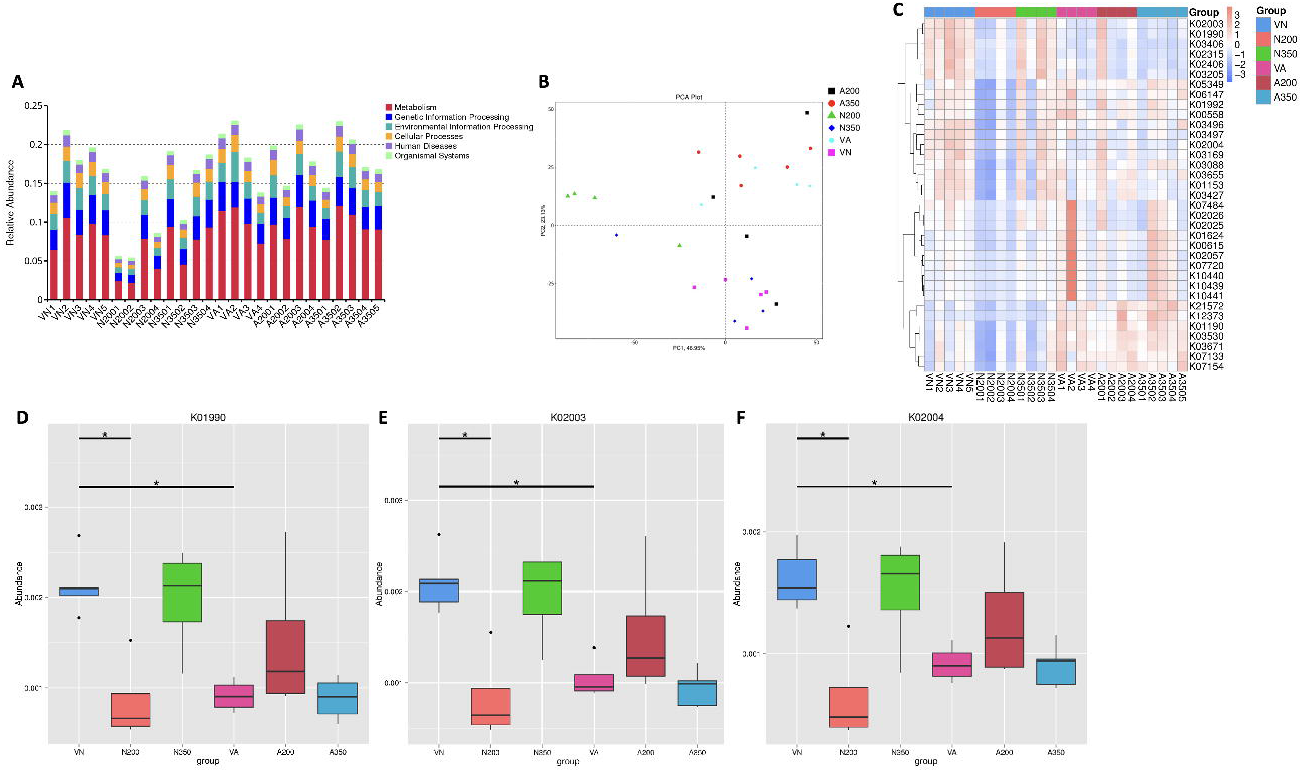
**A:** Relative abundance histogram of functional annotations on level 1 based on the KEGG database. **B:** PCA diagram of the significantly different functions. The abscissa represents the first principal component, and the percentage represents the contribution of the first principal component to the sample difference; the ordinate represents the second principal component, and the percentage represents the contribution of the second principal component to the sample difference. Each point in the graph represents a sample, and samples in the same group are represented by the same color. **C:** Abundance clustering heatmap of significantly different functions based on the KEGG pathway: horizontal is sample information; vertical is functional annotation information; a functional clustering tree is shown on the left side of the figure; the value corresponding to the middle heatmap is each row, and Z-values obtained after normalization of functional relative abundance. **D-F:** Box plot of significantly different functions. The horizontal axis is the sample grouping, and the vertical axis is the relative abundance of the corresponding function. The horizontal line represents two groups with significant differences (*q-value < 0.05; **q-value < 0.01).

### 6. DOR triggers metabolic reprogramming of CD8^+^ T-cells and enhances tumor infiltration of T-cells in a gut microbiota-dependent fashion

Converging lines of evidence suggest that some commensal bacteria and their metabolites modify the host immune system (29–31). This led us to further analyze spleen T-cell functions. To identify alterations caused by DOR, we analyzed splenic CD8^+^ T-cells from four independent cohorts of mice using a nontargeted metabolomics approach to explore metabolic reprogramming following DOR administration. A total of 489 metabolites were detected. It is noteworthy that more than 22.69% (111 out of 489) of significantly altered metabolites in splenic CD8^+^ T-cells were increased in the 200 N group of mice compared with V N controls (Table S3), while a different pattern was detected in the 200 A group compared with V A mice (6.96%, 34 out of 489 metabolites) (Table S4). PCA analyses showed a different clustering pattern between normal and ABX samples. PC scores demonstrated that 200 N mice clustered more consistently and were obviously separated from their controls (V N) (Figure S6A, up). In contrast, 200 A mice clustered separately but with a partial overlap with their controls (V A) (Figure S6A, down). These results indicate that the distribution of 200 N metabolites was significantly different and driven by the phylum Firmicutes. Accordingly, the abundance of several classes of metabolites changed noticeably in the cohorts. Out of 111 metabolites modified in 200 N relative to V N mice, 39 were significantly upregulated and 72 were significantly downregulated (Figure S6B, up, Table S3). Conversely, of the 34 metabolites altered in 200 A compared to V A mice, 14 were significantly enriched and 20 were significantly downregulated. (Figure S6B, down, Table S4).

Hierarchical clustering analysis (HCA) of the animals studied revealed distinct metabolic profiles in 200 N compared to V N mice (Figure S6C). Moreover, the metabolite profiles in 200 A and V A mice were also largely different, with only a small number of metabolites exhibiting a similarly downregulated pattern in the two groups (Figure S6D). We then attempted to understand which metabolic pathways were representative of a specific feature of splenic CD8^+^ T-cells metabolism following DOR administration. Remarkable differences were in fact detected between the 200 N and V N groups in the KEGG pathway-based differential abundance (DA) analysis (Figure S7A). Among the metabolic pathways upregulated in the 200 N group, a large number of metabolites involved in carbon metabolism, 2-oxocarboxylic acid metabolism, aminoacyl-tRNA biosynthesis, pantothenate and CoA biosynthesis, mineral absorption, protein digestion and absorption, glyoxylate and dicarboxylate metabolism, central carbon metabolism in cancer, arginine biosynthesis, alanine, aspartate and glutamate metabolism showed high DA scores (DA=1). However, the effects of DOR on KEGG pathways were all abolished by antibiotic administration (Figure S7B), suggesting that DOR activity was largely mediated by the above-mentioned pathways.

Through the comprehensive analysis of the pathways in which differential metabolites are positioned (including enrichment analysis and topology analysis), we selected the crucial pathways that are most closely related to these metabolites. We mapped authoritative metabolite databases such as KEGG and PubChem across differential metabolites. The outcome of the metabolic pathway analysis was represented in a bubble plot; each bubble corresponded to a metabolic pathway, and the size of the bubble signified the level of enrichment. Figure 7A shows a strong enrichment of cyanoamino acid metabolism, alanine, aspartate, and glutamate metabolism, aminoacyl-tRNA biosynthesis, glycine, serine, and threonine metabolism. More detailed information for each significantly changed pathway including the total number of metabolites/pathways, number of hits/pathways, raw p- and -ln (p)-value, etc. is listed in Table S5.

**Figure 7:**
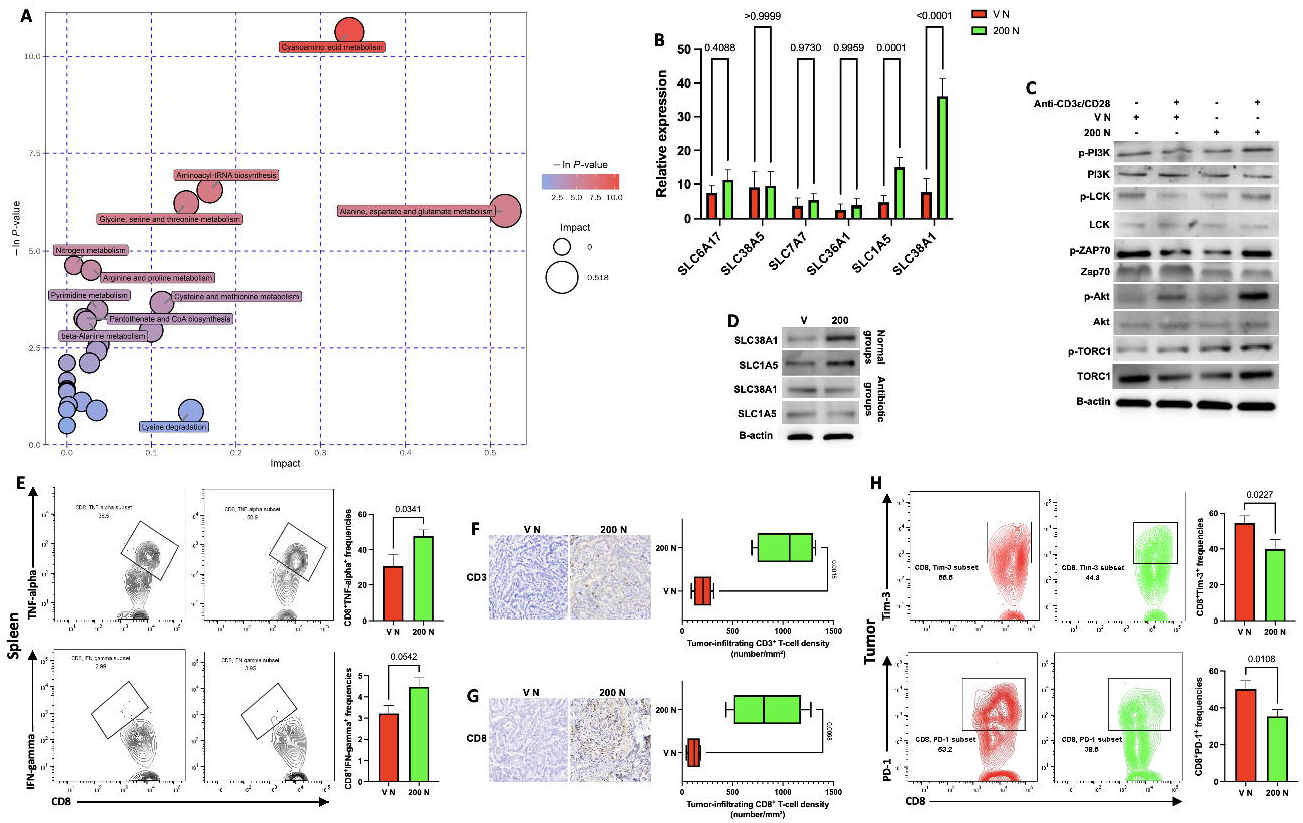
**A:** Metabolic pathway analysis of differential metabolites in groups 200 N *vs* V N. Each bubble in the plot represents a metabolic pathway. The abscissa and the size of the bubble represent the size of the influence factor of the pathway in the topology analysis. The ordinate and the color of the bubble represent the enrichment analysis, whereby a smaller P-value (expressed as -ln P-value) and a darker color represent a more significant enrichment degree. Generally, dark large bubbles are pathways with higher degree of enrichment. **B:** Relative mRNA levels of potential transporters for DOR as well as for amino acid uptake and exchange in CD8^+^ T-cells of groups V N and 200 N were analyzed by qRT-PCR. **C:** DOR promotes T-cell activation through activation of pathways downstream of the TCR. Immunoblot of PI3K, p-PI3K, LCK, p-LCK, Zap70, p-Zap70, Akt, p-Akt, TORC1, and p-TORC1 in CD8^+^ T-cells from mice treated with DOR upon activation with anti-CD3ε/anti-CD28 as indicated. **D:** Western blot showing SLC38A1 and SLC1A5 expression in naive CD8^+^ T-cells activated *ex vivo* from V N and 200 N as well as V A and 200 A. **E:** Representative flow cytometry plots of CD8^+^ T-cells from groups V N and 200 N. Shown are the frequencies of cells expressing TNF-α (E, up) and IFN-γ (E, down). All data are the mean ± SEM of n = 3 independent determinations. **F and G:** IHC staining of CD3^+^ (F) and CD8^+^ (G) TILs 25 days following tumor inoculation in tumor samples from control and DOR-treated mice. **H:** Representative flow cytometry plots of CD8^+^ T-cells from groups V N and 200 N. Shown are the frequencies of cells expressing Tim-3 (H, up) and PD-1 (H, down). All data are the mean ± SEM of n = 3 independent determinations.

Amino acids serve as both an energy source as well as a substrate for protein and nucleic acid biosynthesis in the course of T-cell activation, differentiation, and function. Accordingly, we hypothesized that amino acids transporters reshape amino acid metabolism and, consequently, T-cell function upon DOR administration. To this end, we evaluated the expression patterns of 6 solute carrier (SLC) family transporters in CD8^+^ T-cells. In particular, upon CD8^+^ T-cell activation, the mRNA levels of transporters SLC38A1 and SLC1A5 genes were significantly upregulated in the 200 N group (Figure 7B). From a mechanistic point of view, activation of the pathways downstream of TCR signals and transcription factors containing the mammalian target of rapamycin (mTOR) (Figure 7C) was paralleled by an upregulation of SLC family amino acid transporters in T-cells from DOR-treated mice (Figure 7D), thereby increasing amino acid uptake and further supporting splenic CD4^+^/CD8^+^ T-cell activation, proliferation, and differentiation (Figure 7E and Figure S8A-E). In contrast, amino acids metabolic pathways in 200 A compared to V A mice were significantly reduced and only a high content of purine metabolism and pyrimidine metabolism was seen (Figure S8F). Detailed results of the pathway analyses are shown in Table S6. Intriguingly, depletion of gut microbiota following ABX administration abolished the effect of DOR on expression of both transporters (Figure S8G and Figure 7D), and on the percentages of T-cells expressing activation markers in comparison with control groups (Figure S8H), re-emphasizing the dependence of DOR on the gut microbiota in metabolic reprogramming of CD8^+^ T-cells. To assess whether these changes could be attributed to the development of antitumor immunity, we first analyzed the frequency of tumor infiltrating lymphocytes (TILs) 25 days after tumor inoculation by immunohistochemistry (IHC) with CD3- and CD8-specific antibodies. Compared to control mice, tumors from either group of DOR-treated mice contained higher amounts (number/mm^2^) of CD3^+^ (Figure 7F) and CD8^+^ T-cells (Figure 7G). While, compared to control mice (V A group), tumors from 200 A mice group harbored a lower number/mm^2^ of CD3^+^ and CD8^+^ T-cells (Figure S8I). Next, we further investigated how DOR-mediated reprogramming of amino acid metabolism in CD8^+^ T-cells regulates an exhausted phenotype in the tumor microenvironment. Accordingly, we looked at inhibitory receptors typically expressed by exhausted T-cells by flow cytometry. The data showed reduced numbers of Tim-3^+^CD8^+^ T-cells (Figure 7H, up) and PD-1^+^CD8^+^ T-cells (Figure 7H, down) within 200 N tumors compared to V N tumors, suggesting that activated CD8^+^ T-cells in the tumor microenvironment may facilitate the elimination of tumor cells.

## Discussion

The beneficial effect of commensal microbiota on drug metabolism to enhance CD8^+^ T-cell immunity and increase chemotherapy efficacy has gained substantial attention these last years (32–35). However, the contribution of DOR, a promising anticancer compound, to drive T-cell activation and function against solid tumors remains largely unknown. In this study, using subcutaneously implanted tumors, we benefit from a unique model in which two distinct microbial communities in WT B6 mice are related to diverse disease outcomes. We provide new insight into DOR-mediated host-microbe metabolic crosstalk, which enhances CD8^+^ T-cell activation and infiltration into the tumor microenvironment and is crucial for boosting antitumor immunity against melanoma cells and consequently inhibiting tumor growth in mouse preclinical models. This impact is largely dependent on alterations in the composition of the gut microbiota, as evidenced by the failure of DOR to stimulate antitumor immunity in mice with a dysbiotic microbial community. Our findings are consistent with previous reports indicating that enrichment in the abundance of specific gut microbiota taxa or antibiotic treatment may also inhibit and abolish the effectiveness of immune check-point therapy (36–38). We show that administration of DOR in the presence of a healthy gut microbiota stimulates specific changes in the composition of the fecal microbiota, resulting in translocation of phylum Firmicutes from the gut to MLN/spleen and CD8^+^ T-cell activation. In support of this notion, Viaud *et al*. emphasized that a chemotherapy drug such as cyclophosphamide (CTX) modifies the composition of the small intestine microbiota and induces the translocation of phylum Firmicutes to secondary lymphoid organs, stimulating the generation of a particular subset of “pathogenic” T helper 17 (pTh17) cells and memory Th1 immune responses (39). In particular, Daillère and co-workers found that two Firmicutes species, *Enterococcus hirae* and *Barnesiella intestinihominis*, can increase the intratumoral CD8^+^ T-cell/Treg ratio and the infiltration of IFN-γ-expressing γδ-T-cells during CTX therapy (40). We found that metabolic reprogramming in CD8^+^ T-cells triggered by Firmicutes species increased TNF-α and especially IFN-γ expression in secondary lymphoid organs and tumor infiltration of activated CD8^+^ T-cells, limited T-cell exhaustion, and ultimately decreased tumor burden following DOR administration. We showed that eight bacterial species of the *Clostridia* class were essential and sufficient to enhance cancer-specific effector and memory CD4^+^ and CD8^+^ T-cell function, thus compensating for the limited efficacy of DOR following antibiotics administration. Our findings stressing the importance contribution of the gut microbiota to T-cell responses are in line with previous literature on cancer immunotherapy. For example, a very recent elegant study by Smith and colleagues has shown an association between the therapeutic response to CAR T-cell administration and enrichment of members of the *Clostridia* class. Conversely, higher toxicity and lower survival were seen following exposure to wide-spectrum antibiotics prior to CAR T-cell infusion in patients suffering from B-cell malignancies (41).

In addition, mice receiving ABX also showed reduced body weights relative to untreated mouse groups. This effect was dependent on microbial colonization, as control V N mice exhibited body weights comparable to those of 200 N and 350 N mice, while V A mice displayed significantly delayed body weight gain after the induction of gut dysbiosis by oral antibiotics. Taken together, these results imply a fundamental role for gut microbiota, regardless of DOR administration, in promoting host fitness by stimulating optimal functioning of the “*super organ*” that includes the intestine and microbiota. Therefore, administration of the microbial species identified in this study along with DOR may favor the antitumor effects of DOR and tumor eradication in mice. Further studies are required to fully understand the mechanisms accounting for this interaction.

Considering that microbial metabolites and their derivatives may have immunomodulatory properties (42–45), we then hypothesized that the interaction between DOR and CD8^+^ T-cells involves microbiota derived-metabolites. Our data also showed that specific bacterial metabolites may contribute to providing transporters for DOR absorption and metabolic reprogramming in CD8^+^ T-cells, which resulted in increased production of selected cytokines in MLN and spleen. Conversely, the disruption of Firmicutes communities in antibiotics-treated mice could result in DOR accumulation and toxic effects and ultimately reduce the effectiveness of DOR therapy, increase tumorigenesis, and decrease survival. These data suggest that phylum Firmicutes could alleviate the severity of DOR induced toxicity and, in turn, mitigate T-cell exhaustion to enhance antitumor immunity *via* CD8^+^ T-cell-dependent mechanisms. Furthermore, microbial communities in hosts treated with antibiotics showed reduced diversity and coverage of metabolic pathways, associated with down-regulated CD8^+^ T-cell responses following DOR administration, highlighting the importance of microbial communities in DOR uptake and the regulation of antitumor immunity by DOR. This reduced coverage is probably due to the cumulative impacts of subtle alterations in Firmicutes abundance in the gut microbiota of mice that received 5 days of antibiotic. These overall data suggest that DOR shapes microbial consortia by adjusting the abundance of a few keystone bacteria. Our results are in general agreement with former findings reporting that commensal bacteria regulate the anticancer immune effects of immunotherapy and chemotherapy (37, 39, 46), and that their metabolites, such as butyrate, may enhance the efficacy of chemotherapy with oxaliplatin through modification of CD8^+^ T-cell immunity in the tumor microenvironment (47). Further, the data described herein document that a selective reduction of amino acid metabolic pathways takes place in mice receiving antibiotics, and that gut bacteria, either by regulating host responses or *via* direct metabolic activity, are mandatory for metabolic reprogramming. In support of this notion, the assessment of potential functional roles of gut microbial communities in antibiotic-naïve, normal mouse groups, especially 200 N mice, documented the reduced aggregation of carbohydrate metabolism, amino acid metabolism, translation, and membrane transport in the gut, emphasizing that the translocation of specific gut bacteria provided beneficial effects for DOR metabolization and the development of antitumor immunity. Our findings are in line with Luu *et al*., who demonstrated that microbial short-chain fatty acids (SCFAs), butyrate and pentanoate, generate enhanced antitumor cytotoxic T-lymphocyte (CTL) and CAR T-cell responses through metabolic and epigenetic reprogramming, leading to enhanced production of effector molecules such as TNF-α, CD25 and IFN-γ, and significantly increasing the activity of tumor antigen-specific CTLs and receptor tyrosine kinase like orphan receptor 1 (ROR1)-targeting CAR T-cells in pancreatic cancer and syngeneic murine melanoma models (48).

Our metabolome analysis revealed a remarkable impact of DOR on metabolic pathways linked to CD8^+^ T-cell activation, with extensive alterations in the metabolism of amino acids such as cyanoamino acid, alanine, aspartate, glutamate, glycine, serine and threonine and also aminoacyl-tRNA biosynthesis. An intriguing recent observation has indicated that amino acids, their transporters, and metabolites contribute to the modulation of immune responses (49). Accordingly, previous research studies clearly showed that reprogramming amino acid metabolism has important consequences in the development of tumor immunity and the response to therapy (50, 51). Since the aforementioned amino acid metabolic pathways, which including essential and non-essential amino acids, were up-taken by the CD8^+^ T-cells of 200 N mice, we hypothesized that their diminished load in mice receiving ABX could inhibit the metabolic shift (due to inaccessible amino acid residues) to anabolic metabolism required for T-cell effector functions and expansion and thus impair T-cell proliferation and functions in these animals.

One possible limitation of our study is that it was focused on two mouse models implanted subcutaneously with melanoma cell lines. In fact, extending these observations to different animal models might provide an opportunity to monitor the impact of DOR administration at various phases of tumor progression and in the relevant microenvironment. In addition, given the complex nature of human tumors, studies using patient-derived xenografts are needed to substantiate our observations. The results of such studies may contribute to opening a new window on the use of gut microbiota and its metabolites in cancer chemotherapy.

## Conclusion

In conclusion, the results of this study further our understanding of the antitumor effect of DOR, which, as we show, critically depends on the gut microbiota and its metabolites, representing a link between gut microbes, DOR, and T-cell immunity. In experiments tuning down immune responses, we recognized that in the presence of a normal gut microbiota, DOR can cause bacterial translocation from the gut to the MLN/spleen and promote antitumor immunity through CD8^+^ T-cell activation. In contrast, in mice with ABX-induced dysbiosis, the antitumor effect of DOR would be hindered, and tumorigenesis would be increased due to the absence and reduced diversity of specific microbiota species, in particular the phylum Firmicutes. In addition, increased accumulation of DOR in dysbiotic mice would worsen its toxic effects. Altogether, these observations highlight the importance of the microbial consortium in regulating the antitumor effect of DOR. Therefore, it is anticipated that modification of specific bacterial strains and metabolites along with DOR administration will not only improve our understanding of these interactions but will also increase its therapeutic efficacy and contribute to advancing the development of innovative therapeutic approaches that could be screened across different species and tested for their clinical potential in cancer therapy. Future research is awaited to combine selected gut microbiota species with DOR to activate CD8^+^ T-cells and enhance their antitumor functions in relevant preclinical and clinical models.

## Materials and Methods

### Mice

Wild type C57BL/6 female mice, GF C57BL/6 female mice, and C57BL/6 female mice expressing the congenic markers CD45.1/CD45.2 were purchased from Shanghai SLAC Laboratory Animal Co. Ltd (Shanghai, China) and housed under SPF conditions in gnotobiotic facility in the Animal facility at the Central Laboratory, Clinical Medicine Scientific and Technical Innovation Park (Shanghai Tenth People’s Hospital, Tongji University, Shanghai, China). Mice were used between 6 and 8 weeks of age and were kept under a 12:12-h light-dark (LD) cycle at 22-26°C and were fed with sterile pellet food and water ad libitum. Mouse selection for experiments was not strictly randomized or blinded. GF status was consistently confirmed by aerobic and anaerobic cultures, Gram stains and qPCR as previously described (52). All animal experiments were approved by the Institutional Animal Care and Use Committee (IACUC) of Tongji University (Shanghai, China) and were performed in accordance with the IACUC guidelines and the National Institutes of Health Guide for the Care and Use of Laboratory (National Institutes of Health Publication No. 86-23, 1985).

### Cell lines, culture conditions, and reagents

B16-F10 cells (syngeneic from C57BL/6J mice) were purchased from the American Type Culture Collection (ATCC). B16-OVA cells were purchased from the Sigma-Aldrich (Saint Luis, MO, USA). Both cell lines along with CD8^+^ T-cells were cultured in Dulbecco’s Modified Eagle’s Medium (DMEM; Gibco, USA) supplemented with 10% fetal bovine serum (FBS), 100 IU mb penicillin/streptomycin, 2 mM L-glutamine, 1 mM sodium pyruvate, and MEM nonessential amino acids (Invitrogen) at 37°C under 5% CO_2_. For T-cell cultures this medium was supplemented with 10 ng mL^−1^ IL-2 (R&D, 402-ML). CD8^+^ T-cells were stimulated with anti-CD3ε (5 μg mL^−1^) and anti-CD28 (1 μg mL^−1^) antibodies for up to 40 h. BCPAP (DSMZ, ACC 273) was purchased from German Collection of Microorganisms and Cell Cultures (DSMZ, Braunschweig, Germany) and cultured in RPMI-1640 medium (Biosharp, BL303A) supplemented with 10% FBS and the same type and concentration of amino acids. All cell lines were free of mycoplasma and were authenticated.

### Animal experiments

For tumor growth experiments, mice were injected s.c. with 10^6^ tumor cells. Tumor size was measured by calipers twice a week for calculation of tumor volume. Chemotherapy was conducted by i.p. injection of doramectin (MCE, HY-17035) (200 μg/kg and 350 μg/kg of body weight) or PBS containing equal amounts of DMSO when tumors reached 35-60 mm^2^. Where indicated, mice were treated with a sterile-filtered ABX consisting of ampicillin (Absin, abs811912-200 mg), streptomycin (Absin, abs814442-50 mg), metronidazole (Absin, abs817037-25 mg), vancomycin (Absin, abs815996-250 mg), neomycin (Absin, abs816418-100 mg), and fluconazole (Absin, abs817209-500 mg), each one separately at a concentration of 1 mg mL^−1^ (or 1 g L^−1^) in drinking water for five days to target Gram-positive, Gram-negative and anaerobic bacteria and to inhibit fungal blooms. ABX-containing water was changed every other day due to the short half-life of some antibiotics. CD4^+^ and CD8^+^-T cells were depleted by i.p. injection of 400 mg anti-mouse Thy1.2 (CD90.2, clone 30H12 from Bio X Cell) or rat IgG2b isotype control on days 3, 6, 11, and 16 following tumor inoculation. The efficacy of depletion was verified by FACS analysis of blood samples.

### Isolation and stimulation of CD8^+^ T-cells

For harvest of organs, mice were killed by carbon dioxide administration. Spleen and lymph nodes were dissociated using the frosted ends of microscope slides into RPMI-1640 containing 5% FBS. Cells were passed through a 40-μm cell strainer and then subjected to red blood cell lysis with ACK lysis buffer. Cells were filtered through 40-μm nylon mesh and resuspended in supplemented RPMI-1640. PP cells were harvested from the antimesenteric side of the small intestine and placed into cold supplemented RPMI-1640. Digested PPs were filtered through a 40-μm cell strainer, rinsed with 10 mL of RPMI-1640 containing 2% FBS, then the supernatant was decanted, and the pellet resuspended in 1 mL of RPMI-1640 containing 2% FBS. Tumors were excised, minced, and digested with 1 mg mL^−1^ collagenase D (Absin, abs47048003-100 mg) and 100 mg mL^−1^ DNase I (Absin, abs47047435) at 37°C for 1 h. Digests were then passed through a 40-μm cell strainer to generate a single-cell suspension. Mouse T-cells were isolated from mouse PP/lymph nodes/spleens and tumors using EasySep Mouse CD8^+^ T Cell Isolation kit (StemCell Technologies). For FACS analysis, cell suspensions were subjected to a percoll gradient (Absin, abs9102-100 mL) for 20 min at 2100 rpm, and frozen at −80°C. Human PBMCs were collected from healthy volunteers under approved protocols of the Shanghai Tenth People’s Hospital Ethics Board, and informed consent was obtained from all the participants. Human T-cells were isolated from PBMCs by Ficoll gradient centrifugation (Ficoll-Paque PLUS, GE Healthcare). CD8^+^ T-cells were plated in the presence of 5 μg mL^−1^ plate-bound anti-CD3ε (clone145-2C11, BioLegend) and 1 μg mL^−1^ soluble anti-CD28 (eBioscience) and incubated for 1.5 h at 37°C and 5% CO_2_. Activation of mouse or human CD8^+^ T-cells was assessed by flow cytometry. To determine the direct *in vitro* effect of doramectin on isolated CD8^+^ T-cells, cells were cultured with increasing doses of doramectin (MCE, HY-17035) or corresponding amounts of DMSO (vehicle control) at 37°C and 5% CO_2_ prior to stimulation with anti-CD3ε and anti-CD28. The OT-I TCR transgenic T-cells expressing the CD45.1 allele were stimulated with 1 μg mL^−1^ OVA_257–264_ peptide (SIINFEKL) (MCE, HY-P1489A) in Freund’s complete adjuvant in the presence of 12.5 U mL^−1^ IL-2.

### Flow Cytometry

The isolated cells were stimulated in complete RPMI-1640 (containing 10 mM HEPES, 1% nonessential amino acids and L-glutamine, 1 mM sodium pyruvate, 10% heat-inactivated FBS, and antibiotics) supplemented with eBioscience™ Cell Stimulation Mixture (500X) (00-4970-93) and eBioscience™ protein transport inhibitor mixture (500X) (00-4980-93) for 4 h before intracellular cytokine staining following the manufacturer’s protocols. Cells were then surface stained, fixed and permeabilized using the Fixation/Permeablization kit (eBioscience, cat# 88-8824-00) following the manufacturer’s protocols. Fluorophore-coupled monoclonal antibodies were used to target the following markers: CD8a (clone 5H10-1), CD8a (clone RPA-T8), CD4 (clone GK1.5), CD25 (clone 3C7), CD62L (clone MEL-14), CD45.1 (clone A20), CD45.2 (clone 104), CD69 (clone H1.2F3), CD44 (clone IM7), TNF-α (clone MP6-XT22), IFN-γ (clone XMG1.2), and Granzyme B (clone QA16A02) (all from BioLegend). Dead cells were excluded using Zombie Aqua™ Fixable Viability kit (BioLegend, cat# 423102). A BD LSRFortessa™ X-20 Cell Analyzer was used for measurement of samples. Cell-associated fluorescence was analyzed using FlowJo (Tree Star, Ashland, OR).

### Western blotting

After activation/stimulation, CD8^+^ T-cells were lysed with RIPA total protein extraction lysis buffer (Bioworld Technology, Inc., cat# BD0031), and lysates subsequently subjected to SDS-PAGE. Proteins were transferred to nitrocellulose membranes (Osmonics Inc., MN, USA). Membranes were blocked and incubated with primary antibodies specific for AKT (9272, 1:1000), phospho-Akt (Ser473) (4060, 1:2000), LAT (45533, 1:1000), phospho-LAT (Tyr255) (54364, 1:1000), LCK (2752, 1:1000), phospho-LCK (Tyr394) (70926, 1:1000), Zap-70 (3165, 1:1000), phospho-Zap-70 (Tyr319) (2717, 1:1000), PI3 Kinase p85 (4292, 1:1000), phospho-PI3 Kinase p85 (Tyr458)/p55 (Tyr199) (4228, 1:1000), S6 ribosomal protein (2217, 1:1000), phospho-S6 ribosomal protein (Ser235/236) (81736, 1:1000), TORC1/CRTC1 (2501, 1:1000), and phospho-TORC1/CRTC1 (Ser151) (PA5-106213, 1:1000), followed by HRP rabbit antimouse IgG (minimal x-reactivity) secondary antibodies. The visualization of the blots was performed using the standard protocol for electrochemiluminescence (ECL; Millipore, Bedford, MA, USA).

### Real-time PCR

Total RNA was extracted using Trizol (Invitrogen) and was transcribed to cDNA by using a high-capacity cDNA reverse transcription kit (Applied Biosystems, CA). Real-time PCR was carried out using Sybr Green RT-PCR kits (Invitrogen) on a 7900HT fast real-time PCR system (Applied Biosystems), according to the manufacturer’s instructions, using the following primer sets: SLC6A17, forward primer: 5’-GCACAAGAGAAACAAGCGCA-3’, reverse primer: 5’-CTGGTGACACACCTCTGACC-3’; SLC38A1, forward primer: 5’-CGTATGCTCGGGTACCACC-3’, reverse primer: 5’-CTTCTTGCTCCTCTTCCCCC-3’; SLC36A1, forward primer: 5’-GACTCCATATGGGCACTCGG-3’, reverse primer: 5’-TGACCCTTGTGCGTAGTGTC-3’; SLC7A7, forward primer: 5’-AGGAAGGAGGAGCTGTCAGA-3’, reverse primer: 5’-TGACCATGGCGAAGAGAAGG-3’; SLC38A5, forward primer: 5’-AGACAGAAAAGGGCAGACGG-3’, reverse primer: 5’-TCCTGCATTTCCATCCCCAC −3’; SLC1A5, forward primer: 5’-TTAGGAGCCTTGTTGCTGGG-3’, reverse primer: 5’-TTGAATGGGGGACAGAACGG-3’; β-actin, forward primer: 5’-GATCAAGATCATTGCTCCTCCTGA-3’, reverse primer: 5’-CAGCTCAGTAACAGTCCGCC-3’. Relative expression of genes in each group of mice was normalized to housekeeping β-actin transcript using the 2^−ΔΔCt^ method.

### Immunohistochemistry (IHC)

Four-mm sections of formalin-fixed, paraffin-embedded (FFPE) tumor tissue were deparaffinized in xylene, rehydrated in graded ethanol, and subjected to antigen retrieval by incubation in citrate buffer. IHC analysis was performed by using rat anti-mouse antibodies against CD3 (clone 17A2, eBioscience) and CD8a (clone 53-6.7, eBioscience). Anti-rat secondary antibody was used for detection of primary antibodies and visualized by SignalStain DAB Substrate kit (Cell Signaling Technology, Danvers, MA). All stained tissues were blindly evaluated by pathologists. The densities of cells expressing CD3 and CD8 were measured by quantifying positively stained cells in five random square areas (1 mm^2^ each) within the tumor. The average total number of cells positive for each marker in the five square areas was expressed in density per mm^2^.

### Stool collection and DNA extraction, library construction, quality control and sequencing

Fresh stool pellets were collected at the specified time points, and the samples were immediately frozen and stored at −80°C. Genomic DNA was extracted from the feces using the QIAamp DNA Stool Mini kit (Qiagen, Hilden, Germany), according to the manufacturer’s instructions. Genomic DNAs were randomly broken into specific short fragments with a length of about 350 bp by Covaris ultrasonic crusher. The obtained fragments were end-repaired, A-tailed and further ligated with Illumina adapter and sequence libraries were constructed according to Illumina’s instruction. The fragments with adapters were PCR amplified, size selected, and purified, and the library was checked with Qubit and real-time PCR for quantification and Agilent 2100 bioanalyzer for size distribution detection. Quantified libraries were pooled and sequenced on Illumina platforms (Illumina PE150), according to effective library concentration and data amount required. In order to ensure the accuracy and reliability of subsequent analysis results, the raw data obtained from the Illumina HiSeq sequencing platform were preprocessed using Readfq (V8, https://github.com/cjfields/readfq) to acquire the clean data for subsequent analysis. The raw Illumina paired-end reads were screened according to the following criteria: (1) reads in which the N base has reached a certain percentage (default length of 10 bp) were removed; (2) reads which shared the overlap above a certain portion with adapter (default length of 15 bp) were removed; and (3) low-quality (≤38) bases at the 30 ends of reads were trimmed. The remaining high-quality clean data were blasted to the host database using Bowtie 2.2.4 software (http://bowtiebio.sourceforge.net/bowtie2/index.shtml) to filter the reads that are of host origin. The clean data were assembled and analyzed by SOAPdenovo software (V2.04, http://soap.genomics.org.cn/soapdenovo.html), then the assembled scaftigs from N connection interrupted to obtain sequence fragments without N. All samples’ clean data were respectively compared to each assembled scaffolds by Bowtie2.2.4 software to acquire the PE reads. All the reads not used in the forward step of all samples were combined and then SOAPdenovo software was used for mixed assembly with the same parameters as single assembly. Fragments shorter than 500 bp in all scaftigs generated from single or mixed assembly were filtered out to take advantage of as many reads as possible, and statistical analysis as well as subsequent gene prediction were performed.

### Gene prediction and abundance analysis

MetaGeneMark software (V2.10, http://topaz.gatech.edu/GeneMark/) was used to predict open reading frames (ORF) and extract the gene sequences of the valid contigs from the assembly and the length information shorter than 100 nt filtered out from the predicted results. Then, CD-HIT software (V4.5.8, http://www.bioinformatics.org/cd-hit) was adopted to obtain a unique non-redundant initial gene catalogue and sequences with 90% coverage and 95% identity were considered for further processing. Using Bowtie2.2.4 software, the clean data of each sample were mapped to the initial gene catalogue to calculate the number of reads of the genes compared in each sample. Finally, the genes whose number of reads was ≤ 2 in each sample were filtered out to achieve the final gene catalogue (unigenes) for subsequent analysis. Based on the abundance information of each gene in each sample in the gene catalogue (the number of mapped reads and the length of gene), basic information statistics including core-pan gene analysis and correlation analysis between samples were performed.

### Taxonomy prediction

Taxonomy prediction was conducted by DIAMOND software (V0.9.9, https://github.com/bbuchfink/diamond/) which blasts the unigenes to the sequences of bacteria which are all extracted from the NCBI NR database (Version: 2018-01-02, https://www.ncbi.nlm.nih.gov/). After filtering, the resulting output files were imported into MEGAN as a way of assigning taxonomy to each sequence according to the LCA algorithm and the abundance information of each sample in each taxonomy hierarchy (kingdom, phylum, class, order, family, genus, species), and the number of genes in each sample at each taxonomic level were obtained based on the LCA annotation result and the gene abundance table. The following analysis including Krona analysis, the exhibition of generation situation of relative abundance, the exhibition of abundance cluster heat map, PCA (R ade4 package, version 2.15.3) and NMDS (R vegan package, version 2.15.3) decrease-dimension analyses were performed, which were based on the abundance table of each taxonomic hierarchy. Also, the difference between groups was tested by Anosim analysis (R vegan package, version 2.15.3) and Metastats and LEfSe multivariate statistical analysis were used to look for the different species within groups.

### Common functional database annotations

We adopted DIAMOND software (V0.9.9) to blast unigenes to functional databases including KEGG database (version 2018-01-01, http://www.kegg.jp/kegg/), eggNOG database (version 4.5, http://eggnogdb.embl.de/#/app/home), and CAZy database (version 201801, http://www.cazy.org/) and the alignment result with the highest score (> 60) and E-value < 1e-5 were selected for subsequent analysis. The relative abundance of different functional hierarchy was clustered, and, based on the function annotation result and gene abundance table, the gene number table of each sample in each taxonomy hierarchy was obtained. In addition, the relative abundance profile, the abundance cluster heat map, the decrease-dimension analysis of PCA and NMDS, the Anosim analysis of the difference between/within groups based on functional abundance, comparative analysis of metabolic pathways, and the Metastat and LEfSe analysis of functional difference between groups were performed.

### Metabolite Extraction

CD8^+^ T-cells were isolated from the spleens of naive female mice by negative selection using magnetic beads and activated *in vitro* in the presence of 5 μg mL^−1^ plate-coated anti-CD3ε and 1 μg mL^−1^ soluble anti-CD28 (1.5 h at 37°C and 5% CO_2_). Activated mouse CD8^+^ T-cells were assessed by flow cytometry and subjected to analysis of the metabolic phenotype. Then 1000 μL extract solution (acetonitrile/methanol/water (2:2:1, v/v)) containing isotopically labelled internal standard mixture was added to each sample and the samples were homogenized at 35 Hz for 4 min and sonicated for 5 min in ice-water bath. The homogenization and sonication cycle were repeated 3 times. Then the samples were incubated for 1 h at −40°C and centrifuged at 12000 rpm for 15 min at 4°C. The resulting supernatants were transferred to a fresh glass vial for LC/MS analysis.

### LC-MS/MS Analysis

LC-MS/MS analyses were performed using an UHPLC system (Vanquish, Thermo Fisher Scientific) with a UPLC BEH Amide column (2.1 mm × 100 mm, 1.7 μm) coupled to Q Exactive HFX mass spectrometer (Orbitrap MS, Thermo Fisher Scientific). The mobile phase consisted of 25 mmol L^−1^ ammonium acetate and 25 ammonia hydroxide in water (pH = 9.75) (A) and acetonitrile (B). The auto-sampler temperature was 4°C, and the injection volume was 2 μL. The QE HFX mass spectrometer was used for its ability to acquire MS/MS spectra on information-dependent acquisition (IDA) mode in the control of the acquisition software (Xcalibur, Thermo Fisher Scientific). In this mode, the acquisition software continuously evaluates the full scan MS spectrum. The ESI source conditions were set as following: sheath gas flow rate as 30 Arb, aux gas flow rate as 25 Arb, capillary temperature 350°C, full MS resolution as 60000, MS/MS resolution as 7500, collision energy as 10/30/60 in NCE mode, spray voltage as 3.6 kV (positive) or −3.2 kV (negative), respectively.

### Data preprocessing and annotation

The raw data were converted to the mzXML format using ProteoWizard and processed with an in-house program, which was developed using R and based on XCMS, for peak detection, extraction, alignment, and integration. Then an in-house MS2 database was generated by Shanghai Biotree Biotech Co., Ltd. (Shanghai, China) and applied in metabolite annotation. The cutoff for annotation was set at 0.3.

### Analysis workflow of metabolomics data, metabolic pathways, and statistical analysis

In this study, a total of 12 experimental samples related to 4 groups (V N, 200 N, V A, 200 A) were compared and metabolites were left after relative standard deviation de-noising. Then, the missing values were filled up by half of the minimum value. Also, the total ion current (TIC) normalization method was employed for each sample in this analysis. The final dataset containing the information of peak number, sample name and normalized peak area was imported to SIMCA16.0.2 software package (Sartorius Stedim Data Analytics AB, Umea, Sweden) for multivariate analysis. Data were scaled and logarithmic transformed to minimize the impact of both noise and high variance of the variables. After these transformations, PCA, an unsupervised analysis that reduces the dimension of the data, was carried out to visualize the distribution and the grouping of the samples. 95% confidence interval in the PCA score plot was used as the threshold to identify potential outliers in the dataset. In order to visualize group separation and find significantly changed metabolites, supervised orthogonal projections to latent structures-discriminate analysis (OPLS-DA) was applied. Then, a 7-fold cross validation was performed to calculate the value of R2 and Q2. R2 indicates how well the variation of a variable is explained and Q2 means how well a variable could be predicted. To check the robustness and predictive ability of the OPLS-DA model, 200 times permutations were further conducted. Afterward, the R2 and Q2 intercept values were obtained. The intercept value of Q2 represents the robustness of the model, the risk of overfitting and the reliability of the model, which will be the smaller the better. Furthermore, the value of variable importance in the projection (VIP) of the first principal component in OPLS-DA analysis was obtained. It summarizes the contribution of each variable to the model. The metabolites with VIP > 1 and p < 0.1 (Student’s *t*-test) were considered as significantly changed metabolites. Volcano plots were made using GraphPad Prism (version 9.3.1). Volcano plots can visually display the overall distribution of metabolite differences between groups. HCA was generated using an integrative toolkit-Tbtools (53). Meanwhile, we calculated the Euclidean distance matrix (EDM) for the quantitative value of differential metabolites and clustered the differential metabolites by complete linkage method. Then, we mapped authoritative metabolite databases such as KEGG (http://www.genome.jp/kegg/) through differential metabolites. After obtaining the matching information of differential metabolites, we conducted an enrichment analysis and a topological analysis to find the most critical pathways that are most related to differential metabolites.

### Statistics analyses

The experiments in this study were set up using 3-5 samples or animals per independent group (mouse experiments), condition or repeat. Each experiment was independently repeated. Immunoblot detection, quantitative PCR with reverse transcription, and immunofluorescence staining experiments were performed at least three times. The statistical analyses were performed using GraphPad Prism (version 9.3.1). Unless otherwise noted, all data are shown as mean ± standard error of the mean (SEM). Comparisons were analyzed using a two-way ANOVA or unpaired two-tailed Student’s *t*-test. Differences in Kaplan-Meier survival curves were assessed by the log-rank test.

## Statements and Declarations

## Acknowledgments

We thank Xiaqing Yu and Qiong Luo from the Department of Nuclear Medicine, Shanghai Tenth People’s Hospital, and Yingying Li from School of Pharmacy, East China University of Science and Technology, Shanghai, China for their technical assistance, the Flow Cytometry core (Meier Bao), and the Animal facility at the Central Laboratory, Clinical Medicine Scientific and Technical Innovation Park (Shanghai Tenth People’s Hospital, Tongji University, Shanghai, China). The authors are grateful to Dr. Olivera J. Finn (Department of Immunology, University of Pittsburgh School of Medicine, Pittsburgh, PA, USA) for her critical review of the manuscript and providing excellent support during the writing process. This work was supported by grants from the Natural Science Foundation of Shanghai [21ZR1449600]. Graphical abstract was created with BioRender (https://www.biorender.com).

## Author Contributions

S.T-S. and A.H.M. contributed equally to this work and share joint first co-authorship, conceptualized the study, performed all experiments, designed the analyses, provided the 16S data-processing pipelines, and wrote the manuscript. W.J. processed mouse’ stool samples and provided reagents. V.C. contributed to the writing, interpretation of the results, and review of this study. L.G.B.-H. and F.M. edited the manuscript. Z.L. and D.L. acquired the funding to support this study and supervised the study. Also, all authors read and approved the final manuscript.

## Data Availability Statement (DAS)

Not applicable.

## Ethics approval and consent to participate

Not applicable.

## Consent to participate

Not applicable.

## Consent for publication

Not applicable.

## Figure legends

**Figure S1: A:** Surface expression of CD69, CD25, and CD44 and production of cytokines from mouse OT-I CD8^+^ T-cells activated by OVA peptide in complete medium containing 0.5, 2.5, or 5 μM DOR or DMSO (vehicle control). **B**: OT-I CD8^+^ T-cells were cultured with B16-OVA melanoma cells in the absence or presence of DOR for 40 h. The percentage of annexin V^+^ of co-cultured B16-OVa cells (B, left) and BCPAP cells (B, right) was measured by FACS analysis. **C:** Antibody-mediated depletion of CD4^+^ and CD8^+^ T-cells in V N and 200 N mice. Anti-mouse Thy1.2 or control immunoglobulin G (IgG, 400 mg) were injected 4 times on days 3, 6, 11, and 16 upon tumor inoculation. Tumor volume was assessed on days 7, 11, 15, 19, 23, and 25 following tumor challenge. FACS analysis revealed > 90% depletion of blood CD4^+^ and CD8^+^ T-cells before day 11 (not shown). **D:** Schematic of the *in vivo* priming of T-cells for tumor experiments: C57BL/6 mice were s.c. injected with B16-OVA cells, immunodepleted by wholebody irradiation and transferred with naive OT-I CD8^+^ T-cells pre-activated in medium prior to treatment with DOR or PBS containing corresponding amounts of DMSO (vehicle control). **E and F:** The tumor burden (E) and mouse survival (F) were measured. Asterisks are representative of p-values < 0.05. **G:** Quantification of CD45.1^+^ OT-I CD8^+^ central (CD44^+^CD62L^+^) and effector (CD44^+^CD62L^-^) memory T-cells in the spleens of C57BL/6 mice (CD45.2^+^) treated with or without DOR and injected with B16-OVA melanoma cells. **H:** Western blot analysis of phosphorylated mTORC1 (Ser151) and S6 ribosomal protein (Ser235/236) in *ex vivo* CD8^+^ T-cells stimulated (5 μg mL^−1^ anti-CD3ε and 1 μg mL^−1^ anti-CD28 antibodies) for 15 min following *in vivo* administration of 200 μg/kg of DOR (group 200 N) or PBS control (group V N). **I:** Western blot analysis of S6 and phospho-S6 (Ser235/236) ribosomal protein in mouse naive CD8^+^ T-cells in medium supplemented with the indicated amounts of rapamycin for 12 h.

**Figure S2: A-D and E-H:** Frequencies of CD8^+^CD25^+^ (S2A and S2E), and TNF-α^+^ (S2B and S2F), IFN-γ^+^ (S2C and S2G), and Granzyme B^+^ CD8^+^ T-cells (S2D and S2H) were assessed in the PPs and MLNs in groups V A, 200 A, and 350 A by FACS. **I-P:** DOR induces bacterial translocation in MLN and spleen from either DOR-injected group in the presence of a normal gut microbiota, not in mice with ABX-induced dysbiosis. Colonies were enumerated after 48-h cultivation of MLN and spleen cells from groups V N, 200 N, and 350 N as well as V A, 200 A, and 350 A in aerobic (I, J, M, and N) or anaerobic (K, L, O, and P) conditions.

**Figure S3: A:** Cluster heat map of the relative abundance at the genus level. The horizontal legend shows the sample information and the vertical shows the species information. The clustering tree on the left in the figure is based on the phylum. **B:** Boxplot diagram of the relative abundance of the 12 dominant species in samples from group 200 N and 350 N compared to VN. **C:** Boxplot diagram of the relative abundance of 3 dominant species in 200 N and 350 N mice compared to V N. **D:** Boxplot diagram of the relative abundance of 3 dominant *Akkermansia* species in the V A group compared to the V N group. All data are the mean ± SEM of n = 3 independent determinations.

**Figure S4: A:** eggNOG-based relative abundance histogram of functional annotations on level 1 belonging to the normal and antibiotic groups (V N, 200 N, and 350 N; V A; 200 A, and 350 A). **B:** PCA plot based on functional abundance in eggNOG database. **C:** Functional abundance clustering heatmap. Samples are listed at the bottom of the map, and functional annotations are shown on the right side. The functional clustering tree is shown on the left side of the figure and the value corresponding to the middle of the heat map is Z-value obtained after normalizing the relative abundance of each row of functions.

**Figure S5: A:** CAZy-based relative abundance histogram of functional annotations on level 1 belonging to the normal and antibiotic groups. **B:** PCA plot based on functional abundance in CAZy database. **C:** Abundance clustering heatmap based on significantly different functions according to CAZy database. Legends at the bottom and right side of the map indicate distinct samples and functional annotation information, respectively. The functional clustering tree is shown on the left side of the figure and the value corresponding to the middle of the heatmap is each row Z-values obtained after normalization of functional relative abundance.

**Figure S6: A:** PCA score plot for 200 N in comparison to V N (up) and 200 A in comparison to V A (down). The abscissa PC1 and the ordinate PC2 in the figure show the scores of the first and second principal components, respectively, each scatter point represents a sample, and the color and shape of the scatter point represent different groups. The sample is basically within the 95% confidence interval (Hotelling’s T-squared ellipse). **B:** Volcano plot for group 200 N *vs* V N (up) and 200 A *vs* V A (down). Each point in the volcano plot represents a metabolite. The abscissa represents the fold change of each substance in the group (log_2_ fold change) and the ordinate represents the P-value of the Student’s *t*-test (-log_10_). The size of the scatter points represents the VIP value, for example, the larger the scatter, the greater the VIP value and vice versa. A complete list of significantly altered metabolites is also presented in Tables S3 and S4 besides being pictured in this figure. Significantly upregulated and significantly downregulated metabolites are shown in red and blue, respectively (in both the Figure and the Tables), while non-significantly different metabolites are shown in grey. **C:** Heatmap of HCA for groups 200 N *vs* V N. Individual animals within the experimental groups are listed at the bottom of the map and the differentially expressed metabolites are listed on the right side. The color blocks at different positions represent the relative expression of the metabolites at the corresponding positions, red and blue indicating that the metabolite is expressed at higher and lower levels in their group, respectively. **D:** HCA heatmap between groups 200 A and V A. Individual animals are listed at the bottom of the map and listed on the right side are the differential metabolites compared in the two groups. The color blocks at different positions represent the relative expression of the metabolites at the corresponding positions, red and blue indicating that the metabolite is expressed at higher and lower levels in their group, respectively.

**Figure S7: A and B:** Pathway-based analysis of metabolomic changes between groups 200 N *vs* V N (A) and 200 A *vs* V A (B). The DA score captures the average alterations for all metabolites in a pathway. A score of 1 implies that those metabolites in the pathway increase in 200 groups compared to their respective V groups, and a score of −1 signifies that those metabolites in a pathway decrease. Pathways with no less than three metabolites were considered for DA score calculation.

**Figure S8: A-E:** The influence of amino acid up-take on CD8^+^/CD4^+^ T-cell function. CD8^+^ T-cell (A-B) and CD4^+^ T-cell (C-E) activation, proliferation and differentiation were evaluated by flow cytometry. **F:** Bubble plot of metabolic pathways correlated with differential metabolites in groups 200 A *vs* V A. Each bubble in the plot represents a metabolic pathway. The abscissa and the size of the bubble represent the size of the influence factor of the pathway in the topology analysis. The ordinate and the color of the bubble represent the enrichment analysis. A smaller P-value (expressed as -ln P-value) and a darker color represent a more significant enrichment degree. Generally, dark large bubbles are pathways with higher enrichment degree. **G:** Relative mRNA expression of potential transporters for amino acid uptake in CD8^+^ T-cells isolated from groups V A and 200 A was analyzed by qRT-PCR. **H:** CD8^+^ T-cell as well as CD4^+^ T-cell activation were evaluated from mice administered antibiotics. All data are the mean ± SEM of n = 3 independent determinations. **I:** IHC staining of CD3^+^ and CD8^+^ TILs following tumor inoculation in tumor samples from ABX-treated mice administered DOR (vehicle control).

