## Supplementary figures and images for "Host-microbe interactions mediate doramectin-promoted metabolic reprogramming of CD8^+^ T-cells and amplify antitumor immunity"

### Figure S1.png

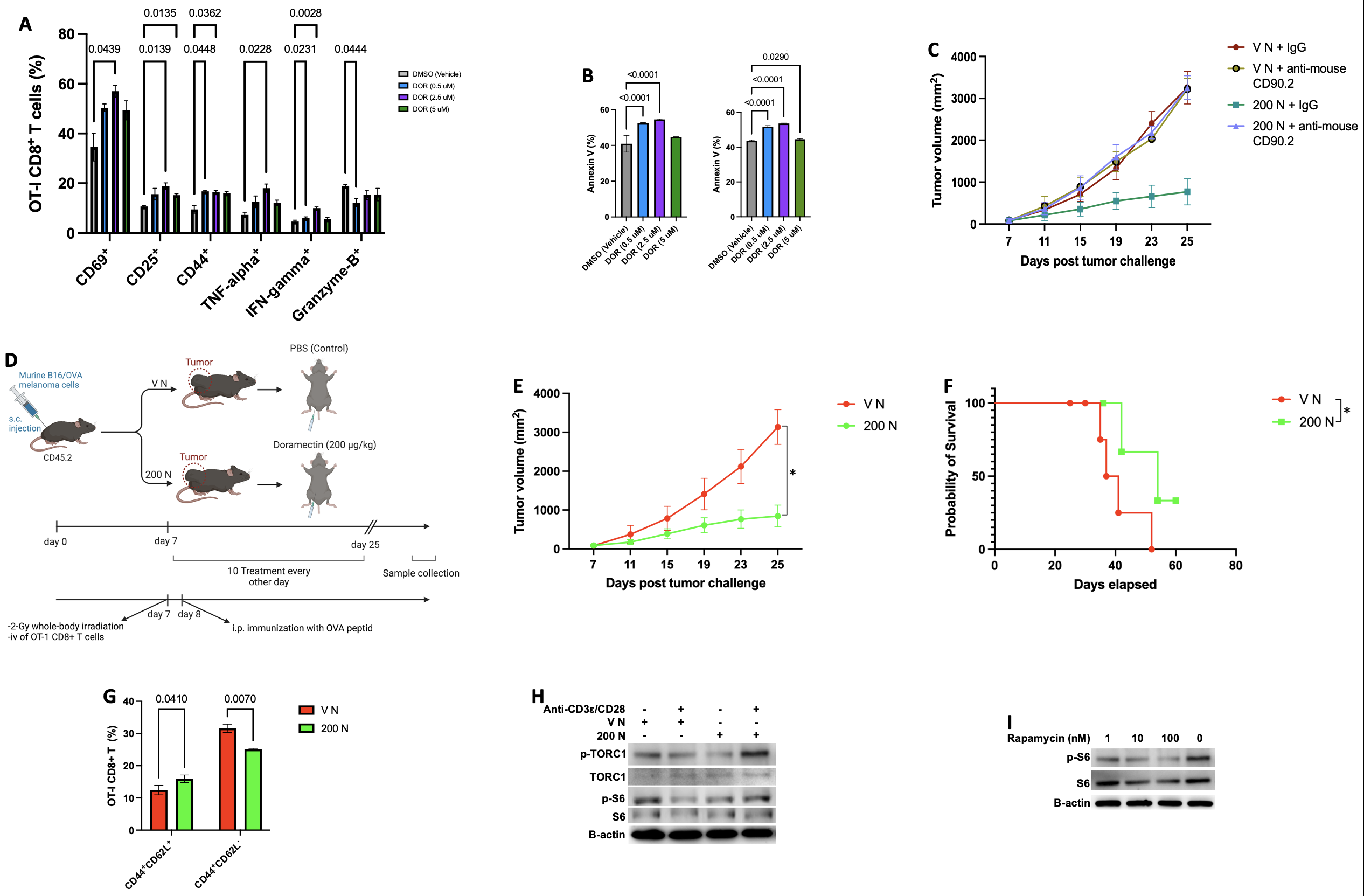

### Figure S2.png

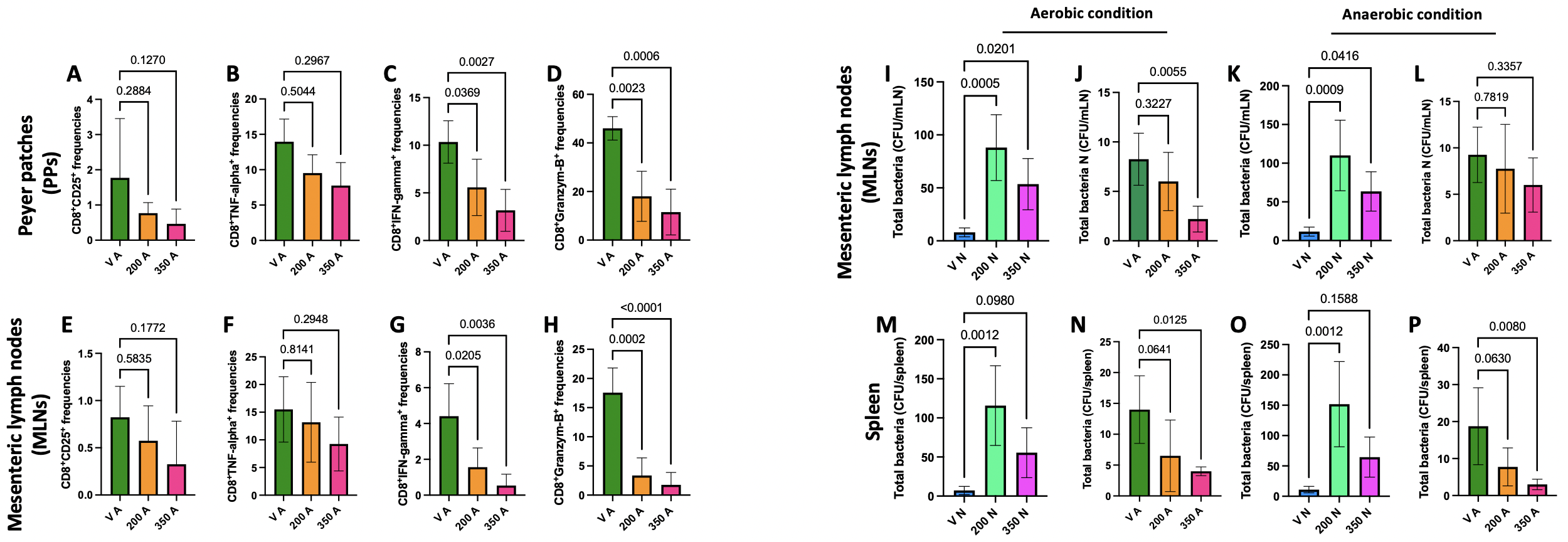

### Figure S3.png

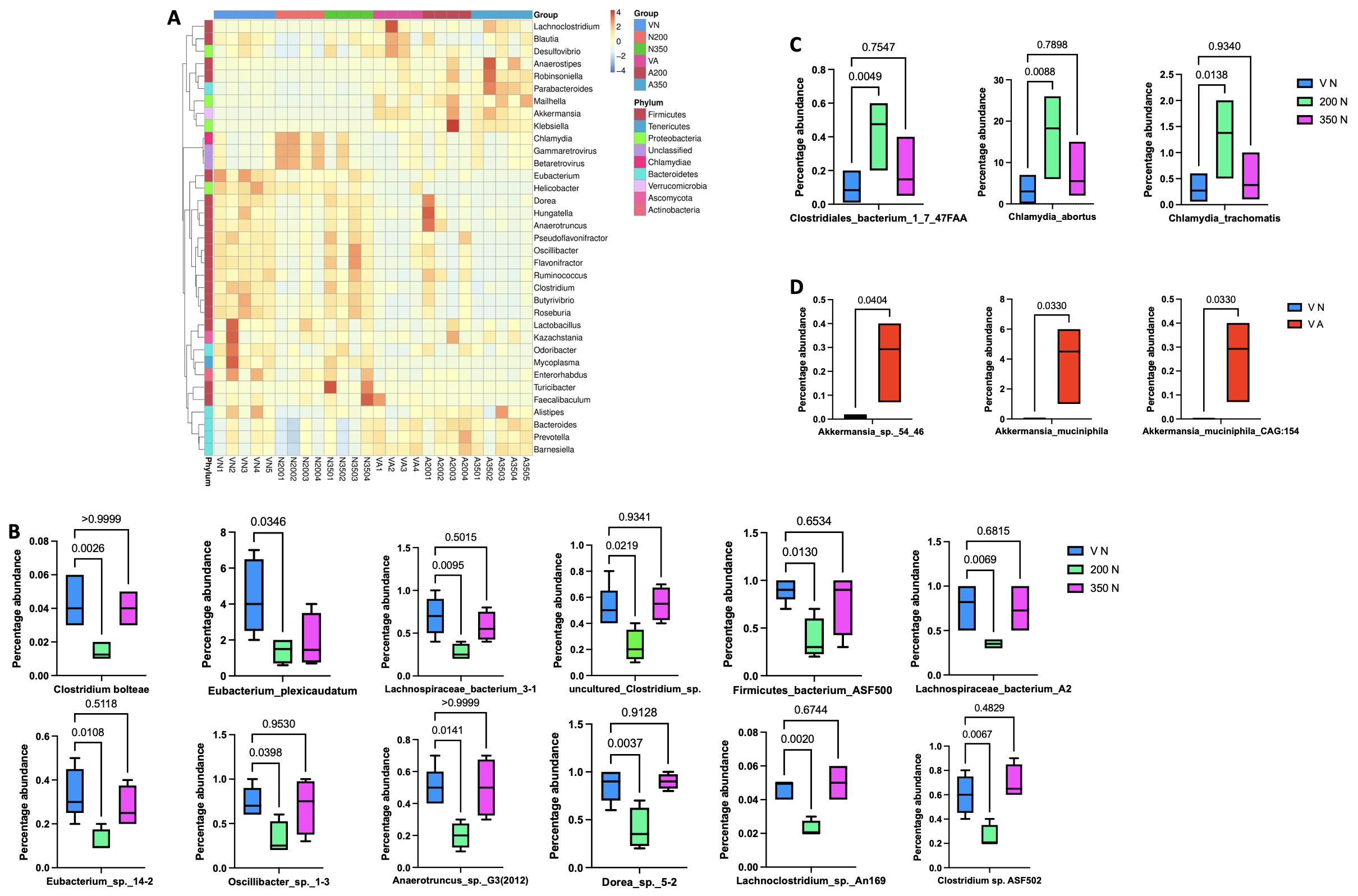

### Figure S4.png

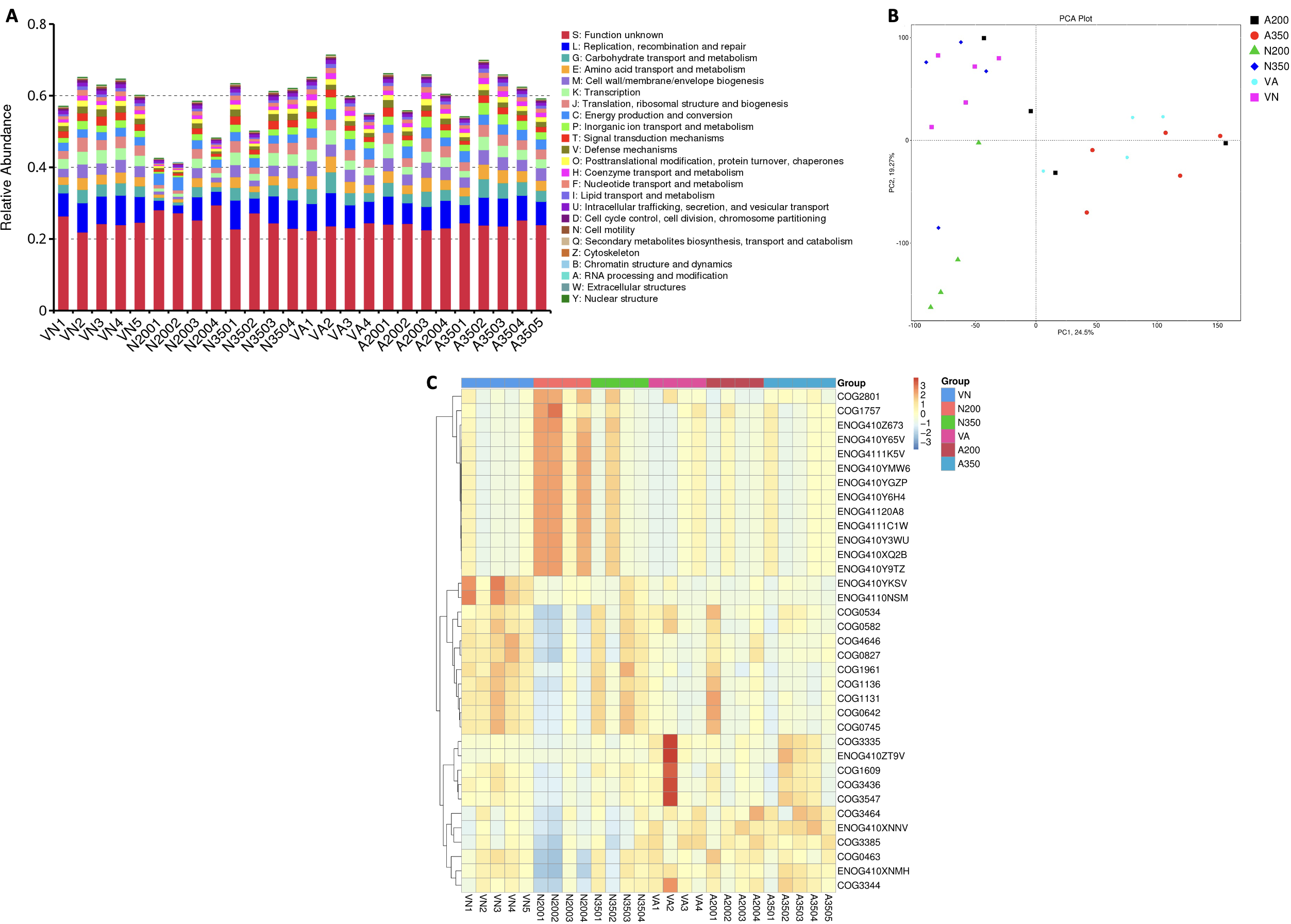

### Figure S5.png

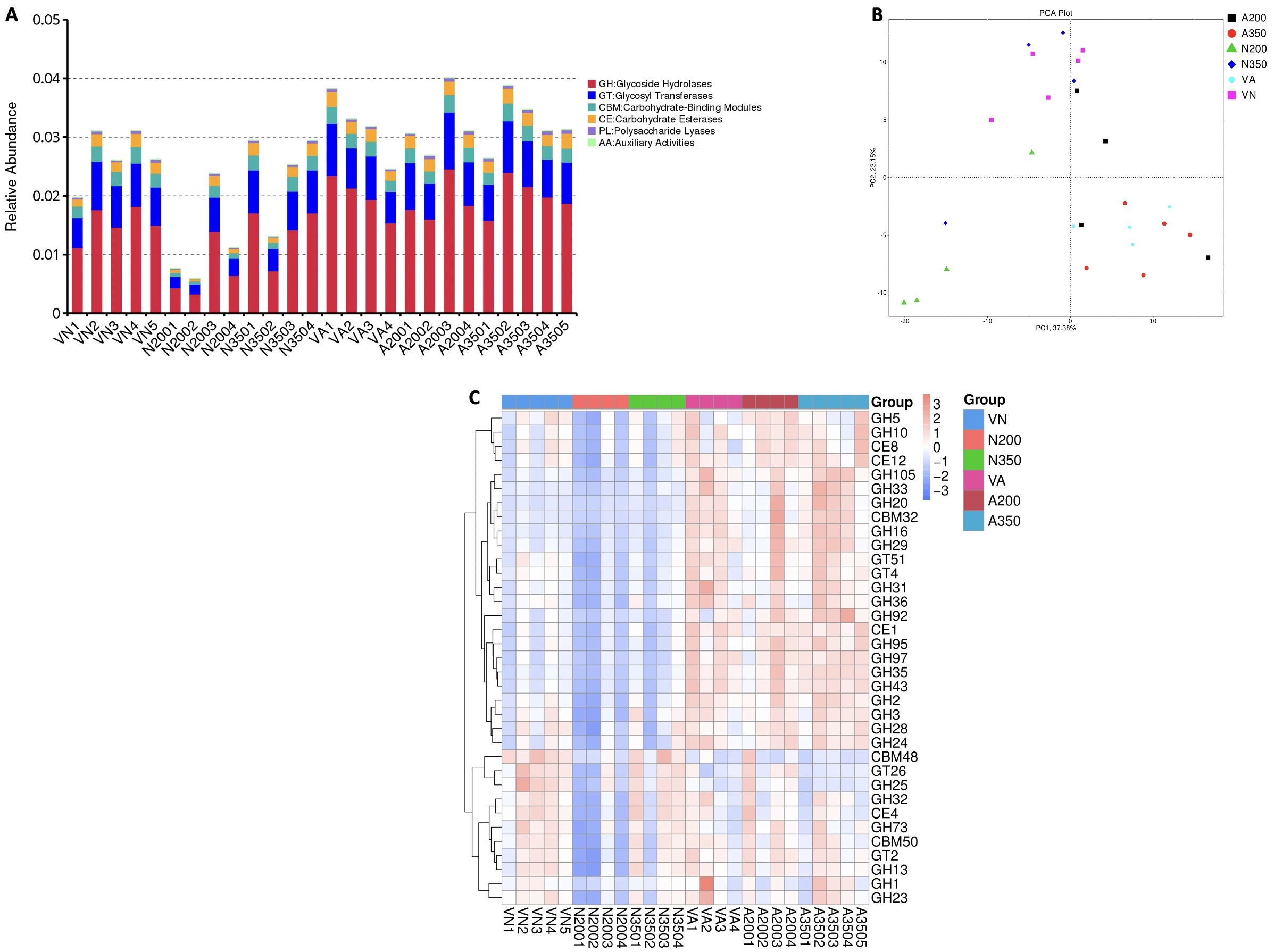

### Figure S6.png

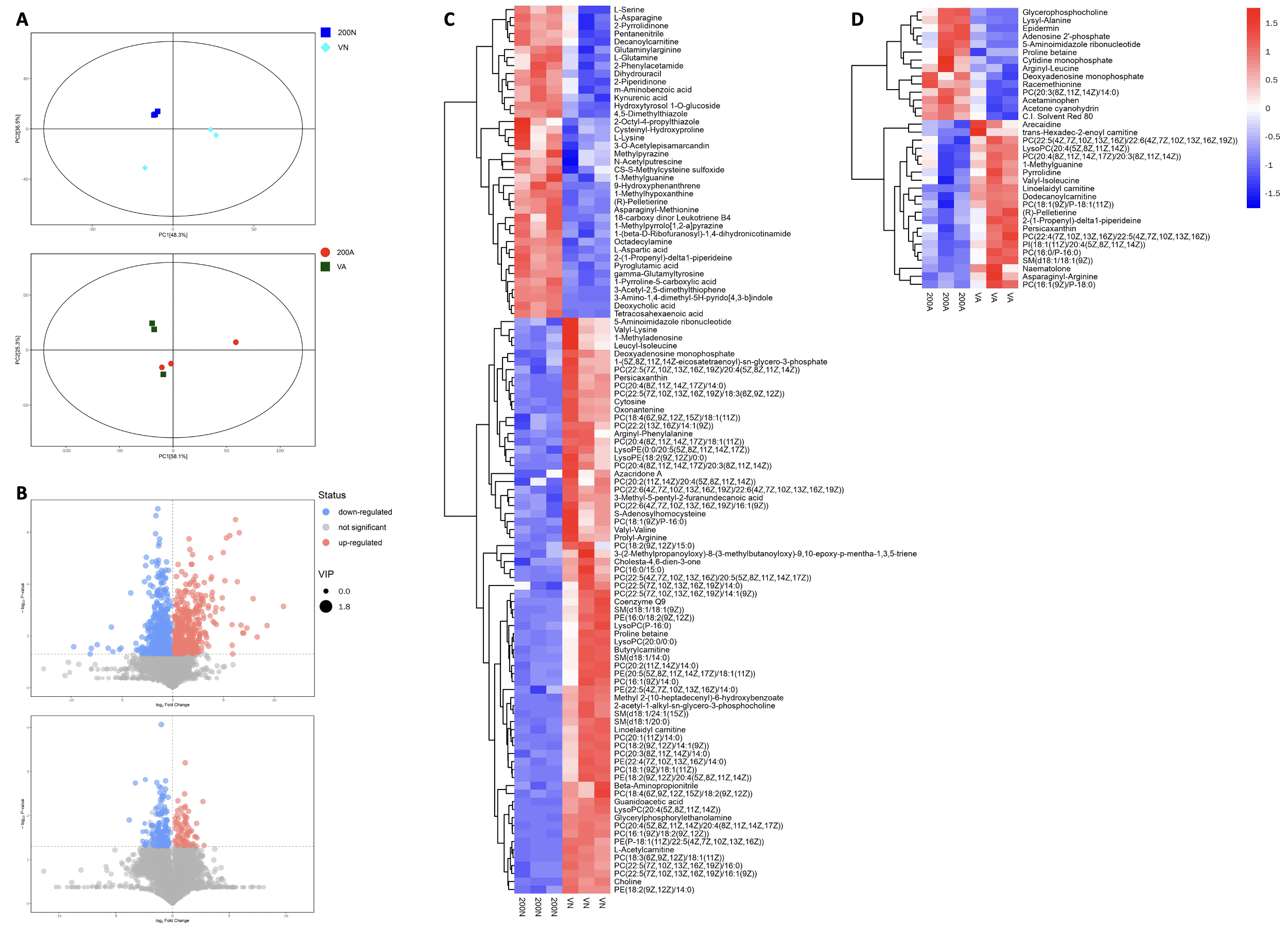

### Figure S7.png

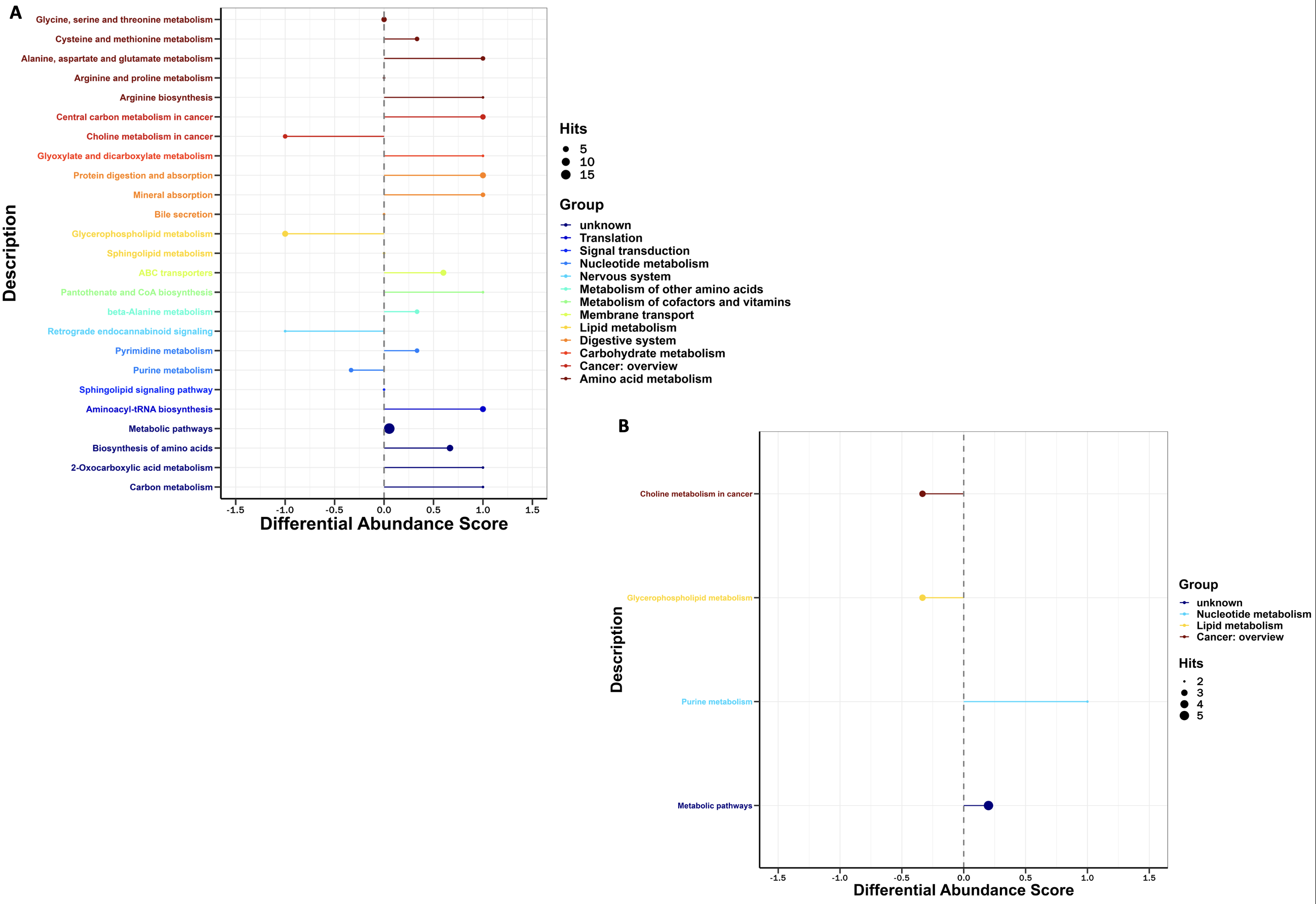

### Figure S8.png

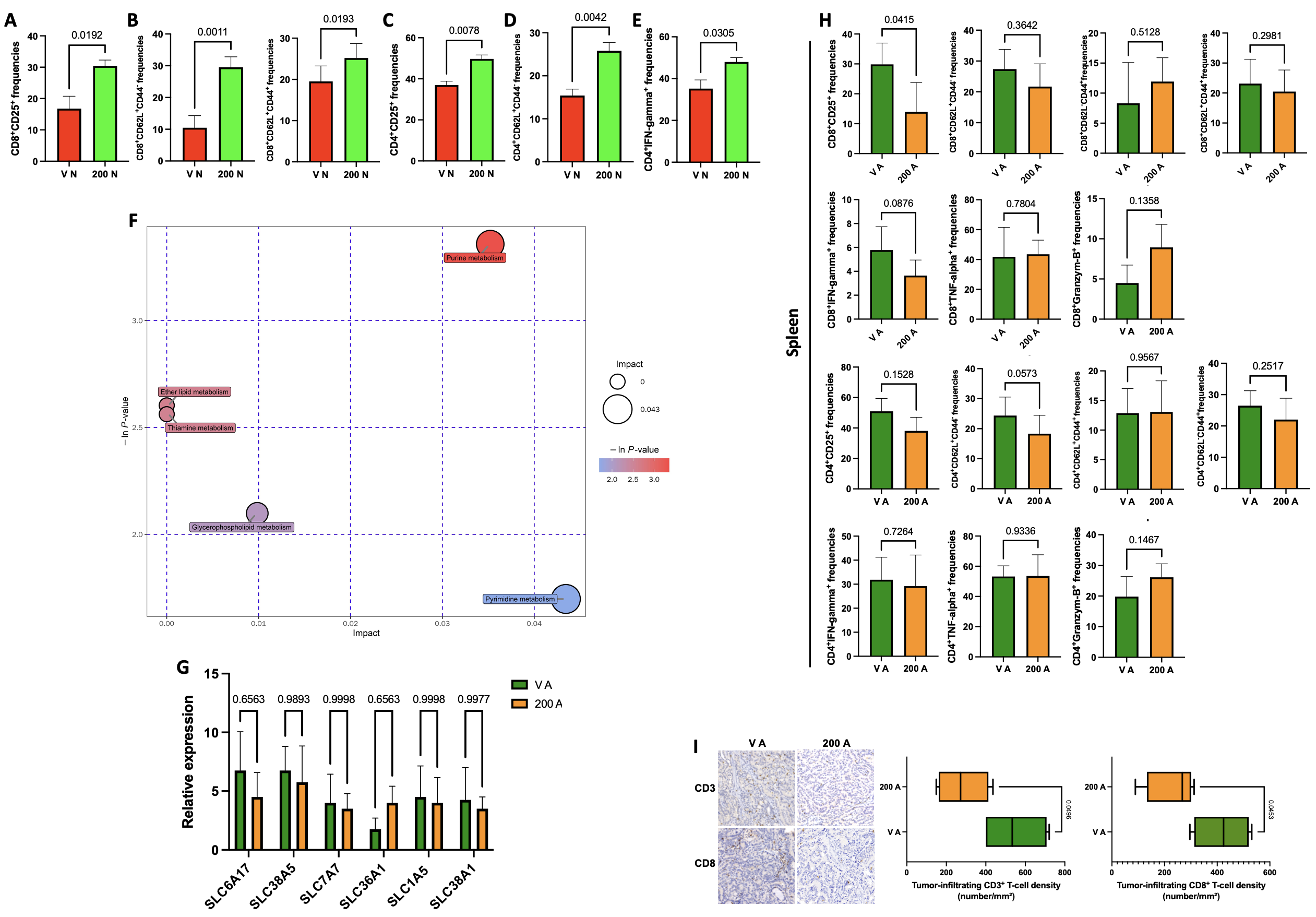
